# An acidic loop in the FHA domain of the yeast meiosis-specific kinase Mek1 interacts with a specific motif in a subset of Mek1 substrates

**DOI:** 10.1101/2024.05.24.595751

**Authors:** Qixuan Weng, Lihong Wan, Geburah C. Straker, Tom. D. Deegan, Bernard P. Duncker, Aaron M. Neiman, Ed Luk, Nancy M. Hollingsworth

## Abstract

The meiosis-specific kinase Mek1 regulates key steps in meiotic recombination in the budding yeast, *Saccharomyces cerevisiae*. *MEK1* limits resection at the double strand break (DSB) ends and is required for preferential strand invasion into homologs, a process known as interhomolog bias. After strand invasion, *MEK1* promotes phosphorylation of the synaptonemal complex protein Zip1 that is necessary for DSB repair mediated by a crossover specific pathway that enables chromosome synapsis. In addition, Mek1 phosphorylation of the meiosis-specific transcription factor, Ndt80, regulates the meiotic recombination checkpoint that prevents exit from pachytene when DSBs are present. Mek1 interacts with Ndt80 through a five amino acid sequence, RPSKR, located between the DNA binding and activation domains of Ndt80. AlphaFold Multimer modeling of a fragment of Ndt80 containing the RPSKR motif and full length Mek1 indicated that RPSKR binds to an acidic loop located in the Mek1 FHA domain, a non-canonical interaction with this motif. A second protein, the 5’-3’ helicase Rrm3, similarly interacts with Mek1 through an RPAKR motif and is an in vitro substrate of Mek1. Genetic analysis using various mutants in the *MEK1* acidic loop validated the AlphaFold model, in that they specifically disrupt two-hybrid interactions with Ndt80 and Rrm3. Phenotypic analyses further showed that the acidic loop mutants are defective in the meiotic recombination checkpoint, and in certain circumstances exhibit more severe phenotypes compared to the *NDT80* mutant with the RPSKR sequence deleted, suggesting that additional, as yet unknown, substrates of Mek1 also bind to Mek1 using an RPXKR motif.

**ARTICLE SUMMARY:** The FHA domain is conserved module best known for creating protein complexes by binding to phosphorylated threonines on target proteins. This work identified a non-canonical mechanism by which the FHA domain of the yeast meiosis-specific kinase Mek1 interacts with two of its substrates, Ndt80 and Rrm3. An acidic loop within the FHA domain binds to RPXKR motifs in Ndt80 and Rrm3. Genetic evidence suggests that this FHA domain acidic loop is required binding to additional Mek1 substrates.

## INTRODUCTION

Meiotic recombination plays a critical role in sexual reproduction as it produces crossovers between homologous chromosomes. These crossovers, combined with sister chromatid cohesion, physically connect homologs to allow their proper segregation at the first meiotic division. After completion of the second meiotic division when sister chromatids separate to opposite poles, haploid genomes are packaged into gametes such as spores in yeast. In the absence of recombination, chromosome mis-segregation at the first meiotic division produces aneuploid gametes, leading to infertility and birth defects such as Trisomy 21 (Hassold and Hunt 2001; Hassold *et al*. 2007).

Recombination is initiated by the formation of double strand breaks (DSBs) at preferred regions of the genome called hotspots (Keeney *et al*. 2014). Resection of the 5’ ends of each break creates 3’ single stranded tails that are bound by the meiosis-specific recombinase Dmc1, as well as Rad51, to form nucleoprotein filaments (Brown and Bishop 2014). These presynaptic filaments search the genome for homology and mediate strand invasion to form displacement or D-loops (Hunter 2007). After extension of the invading strand by DNA replication, D-loops may be disassembled and the breaks repaired by synthesis-dependent strand annealing to produce noncrossovers (Allers and Lichten 2001; Hunter 2007; Mcmahill *et al*. 2007). Alternatively, the 3’ end on the other side of the break can anneal to the displaced strand from the donor duplex to create double Holliday junction intermediates (Schwacha and Kleckner 1995). In budding yeast, a group of proteins collectively referred to as ZMM proteins protects D-loops from disassembly and generates double Holliday junctions that are preferentially resolved to form crossovers (Borner *et al*. 2004; De muyt *et al*. 2012; Zakharyevich *et al*. 2012; Pyatnitskaya *et al*. 2019). In addition, the intermediates generated by the ZMM pathway are required for homologous chromosomes to synapse by formation of synaptonemal complexes (Borner *et al*. 2004).

Approximately 160 DSBs are generated during yeast meiosis to produce around 90 crossovers distributed over 16 pairs of homologs (Chen *et al*. 2008; Mancera *et al*. 2008; Pan *et al*. 2011). Because packaging a broken chromosome into a spore would be lethal, the repair of meiotic DSBs is highly regulated (Hollingsworth and Gaglione 2019). Mek1/Mre11 (henceforth Mek1) is a meiosis-specific kinase that is activated in response to meiotic DSBs (Rockmill and Roeder 1991; Leem and Ogawa 1992; Carballo *et al*. 2008). The DNA damage checkpoint kinases, Tel1 and Mec1, are recruited to DSBs where they phosphorylate the meiosis-specific axial element protein, Hop1 (Carballo *et al*. 2008; Ho and Burgess 2011). Mek1 binds to phospho-Hop1 through its forkhead-associated (FHA) domain and the kinase is activated by autophosphorylation in *trans* (Niu *et al*. 2007; Carballo *et al*. 2008; Chuang *et al*. 2012). Mek1 kinase activity limits resection of the 5’ ends of the breaks, and biases strand invasion into homologs, as opposed to sister chromatids which are the preferred templates for repair in vegetative cells (Kadyk and Hartwell 1992; Bzymek *et al*. 2010; Kim *et al*. 2010; Grubb and Bishop 2024). The substrate(s) phosphorylated by Mek1 that mediate interhomolog bias are unknown. Mek1 inhibits Rad51 by preventing the recombinase from interacting with its accessory factor, Rad54, through phosphorylation of Rad54 and the meiosis-specific protein, Hed1 (Tsubouchi and Roeder 2006; Busygina *et al*. 2008; Niu *et al*. 2009; Callender *et al*. 2016). In addition, Mek1 is required for the ZMM crossover pathway and chromosome synapsis (Chen *et al*. 2015). Finally, Mek1 is the effector kinase for the meiotic recombination checkpoint (MRC) that prevents cells from entering Meiosis I with unrepaired DSBs by inhibition of the meiosis specific transcription factor Ndt80 (Lydall *et al*. 1996; Xu *et al*. 1997; Chen *et al*. 2018).

In budding yeast, mitotic regulators such as the polo-like kinase Cdc5 and the Clb1 cyclin are degraded upon entry into meiosis (Okaz *et al*. 2012). Expression of these genes later in meiosis is then dependent upon Ndt80 (Chu and Herskowitz 1998). The *NDT80* gene is transcribed in two stages (reviewed in (Winter 2012). First, expression mediated by the transcriptional activator, Ime1, produces a low level of Ndt80 protein. Mek1 binds to Ndt80 through a conserved five amino acid sequence (RPSKR) located between the DNA binding and transcriptional activation domains (Chen *et al*. 2018). Mek1 phosphorylation of the Ndt80 DNA binding domain prevents Ndt80 from activating transcription. When chromosomes synapse due to DSB repair by the ZMM pathway, the bulk of Mek1 and the DSB machinery is removed from chromosomes, resulting in a decrease in Mek1 activity (Subramanian *et al*. 2016; Prugar *et al*. 2017; Mu *et al*. 2020). Ndt80 then activates transcription of its own gene as well as >300 target genes, resulting in Holliday junction resolution to produce crossovers, disassembly of the synaptonemal complex that completely inactivates Mek1 and the repair of any remaining DSBs by Rad51 (Chu *et al*. 1998; Sourirajan and Lichten 2008; Argunhan *et al*. 2017; Prugar *et al*. 2017)hedΔR.

Given that Mek1 is a “master regulator” of meiotic recombination in yeast, discovering its substrates is key to understanding mechanisms that control meiotic DSB repair. The protein-protein interaction between Mek1 and Ndt80 was discovered through a two-hybrid screen that utilized *lexA-MEK1* as bait (Chen *et al*. 2018). In addition to *NDT80*, this screen identified several proteins involved in meiotic recombination as putative Mek1 interactors, including the 3’-5’ DNA helicases *SGS1* [amino acids (aa) 82-614) and *SRS2* (aa 1035-1174), the 5’-3’ DNA helicase, *RRM3* (aa 50-299) and *MMS4* (aa 91-378), a subunit of the structure-specific endonuclease, Mus81-Mms4 (de los Santos *et al*. 2001; Jessop and Lichten 2008; Oh *et al*. 2008; Hunt *et al*. 2019; Ziesel *et al*. 2022).

Rrm3 and Pif1 are related superfamily 1B DNA helicases with 5’-3’ unwinding polarity that are involved in a variety of DNA processes, including maintenance of genome integrity (Boule and Zakian 2006; Byrd and Raney 2017; Muellner and Schmidt 2020). Recent work indicates that these helicases function in meiosis as well. Pif1 localizes to DSBs but is inhibited during Dmc1-mediated recombination by the Mer3-MutLβ complex (Vernekar *et al*. 2021). Deletion of *RRM3* exhibits no obvious meiotic phenotypes in an otherwise wild-type diploid (Ziesel *et al*. 2022). Rad51-mediated strand invasion is constitutively activated during meiosis when the Mek1 phosphorylation site on *RAD54* is mutated and *HED1* is deleted (*RAD54-T132A hed1*Δ, abbreviated *hed*Δ*R*) (Liu *et al*. 2012; Lao *et al*. 2013; Ziesel *et al*. 2022). The *NDT80-mid* (Mek1-Interaction-Defective) allele contains a deletion of the RPSKR Mek1 interaction sequence and is defective in the MRC (Chen *et al*. 2018). Spore viability is significantly reduced when Rad51 is activated in the absence of the MRC (*hed*Δ*R NDT80-mid*), and this viability goes down even further when either *RRM3* or *PIF1* is deleted or meiotically depleted, respectively (Ziesel *et al*. 2022). The *hed*Δ*R NDT80-mid rrm3*Δ combination therefore provides a way to functionally assess *RRM3* mutants in meiosis.

This work identifies Rrm3 as an *in vitro* Mek1 substrate that interacts with the kinase through an RPAKR motif. In addition, it shows that an acidic loop located within the Mek1 FHA domain is specifically required for non-canonical protein-protein interactions with Ndt80 and Rrm3. Alanine substitutions within the acidic loop do not affect Mek1 kinase activation but, in combination with *hed*Δ*R*, exhibit meiotic phenotypes that are more severe than *NDT80-mid*. These results suggest that Mek1 binds to one or more unknown substrates through an RPXKR sequence to mediate additional functions such as interhomolog bias.

## RESULTS

### Mek1 interacts with Rrm3 via a conserved RPAKR motif in the Rrm3 amino terminus

The sequences of proteins identified in a two-hybrid screen using *lexA-MEK1* as bait with a Gal4-activation domain (*GAD*) genomic library were examined for the presence of an RXXKR motif similar to RPSKR in Ndt80 (Chen *et al*. 2018). Whereas *GAD-SGS1^82-614^* and *GAD-MMS4^91-378^* lacked this motif, *GAD-RRM3^51-723^*and *GAD-SRS2^1035-1174^* contained RPAKR [amino acids (aa)185-189] and RKSKR (aa 1072-1076), respectively, raising the possibility that these sequences mediate binding to *lexA-MEK1*. This was not the case for Srs2, however, since *GAD-SRS2^1035-1174^*containing a deletion of the RKSKR sequence still interacted with *lexA-MEK1* (Figure S1A). This result shows that an RXXKR motif alone is not sufficient for binding to Mek1.

The RPAKR motif in Rrm3 is localized to the N-terminus of the protein (aa 1-249) where it is flanked by nine Mek1 consensus phospho-sites (RXXT/S) (Figure 1A) (Suhandynata *et al*. 2016). Three more Mek1 consensus sites are located in the C-terminal part of the protein (aa 561-723). A BLAST search using the Rrm3 protein against a variety of yeast species revealed that the RPAKR sequence within the N-terminal region is conserved. Similar to Ndt80, the KR amino acids are present in all species examined (Figure 1B) (Chen *et al*. 2018). While the original *GAD-RRM3^51-299^* fusion plasmid was lost, interaction with *lexA-MEK1* was reproduced using *GAD-RRM3^51-723^* (Figure 1A, C). *GAD-RRM3^51-723^* containing a deletion of RPAKR or mutation of the positively charged KR to negatively charged aspartates (*KR->DD*) or alanines (*KR->AA*) failed to interact with *lexA-MEK1* (Figure 1C). Immunoblot analysis using α-GAD antibodies to detect GAD-Rrm3 showed that the mutant proteins were stable (Figure 1D).

**Figure 1.**
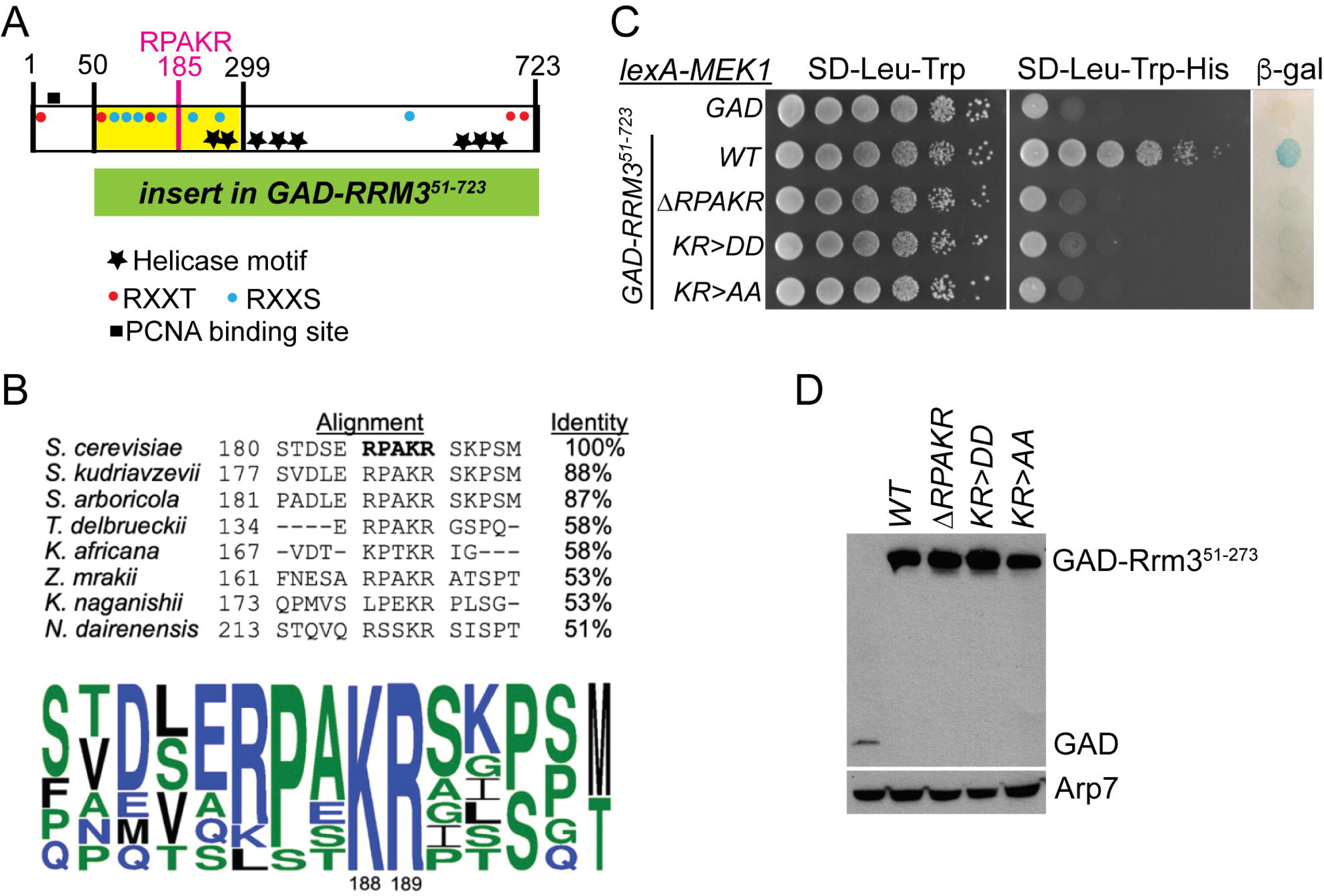
Mek1 interacts with a conserved motif in the Rrm3 N-terminus in the two-hybrid system. (A) Schematic of the Rrm3 protein. The yellow box indicates the 50-299 fragment of Rrm3 found in a two-hybrid screen that used *lexA-MEK1* as bait. The green box indicates the 51-723 Rrm3 fragment used for the two-hybrid analyses in this figure. “RPAKR” indicates the Mek1 interaction sequence, while blue and red dots depict RXXT and RXXS Mek1 consensus phosphorylation sites (T37, T73, S85, S116, S125, T155, S165, S227, S272, S565, T692, T721), respectively. The black box represents a PCNA binding site and stars indicate helicase motifs (Schmidt *et al*. 2002; Bessler and Zakian 2004). Numbers indicate amino acid positions within the Rrm3 protein. (B) (Top) Alignment of a segment of the *S. cerevisiae* Rrm3 protein with other yeast orthologs showing conservation of the RPAKR motif. Numbers on the left indicate the amino positions of the first amino acid in each sequence while numbers on the right indicate percent identity with *S. cerevisiae* Rrm3. (Bottom) A consensus motif based on the alignment generated using WebLogo (Crooks *et al*. 2004). Numbers indicate the amino acid positions of K and R in *S. cerevisiae* Rrm3. (C) Two-hybrid analysis. The L40 strain containing *HIS3* and *lacZ* reporters was transformed with the *lexA-MEK1* plasmid (pTS3), and plasmids containing either *GAD* (pACTII), *GAD-RRM3^51-723^* (pJW1), *GAD-RRM3^51-723^-*Δ*RPAKR* (pJW2), *GAD-RRM3^51-723^-KR>DD* (pJW1-KR>DD) or *GAD-RRM3^51-723^-KR>AA* (pJW1-KR>AA). Ten-fold serial dilutions starting with an equal number of cells were spotted onto either SD-Leu-Trp medium to select for the two plasmids or SD-Leu-Trp-His medium. Growth in the absence of histidine indicates a two-hybrid interaction. Undiluted cells were also spotted onto a filter that was used to measure β-galactosidase produced by transcription of *lacZ* that results in a blue color. (D) Immunoblots of protein extracts from the same cultures used for the spotting assay in C were probed with α-GAD antibodies to detect the different GAD-Rrm3 proteins. α-Arp7 antibodies were used to detect Arp7 as a loading control.

### Mek1 directly phosphorylates Rrm3 *in vitro*

The discovery that Rrm3 binds to Mek1 via an RPAKR motif raised the possibility that it is a Mek1 substrate. This hypothesis was tested using the semi-synthetic epitope system to perform *in vitro* kinase assays using GST-Mek1-as partially purified from meiotic yeast cells (Allen *et al*. 2007; Lo and Hollingsworth 2011). GST-Mek1-as contains an “analog sensitive” mutation that enlarges the ATP binding pocket of the kinase. As a result, only GST-Mek1-as can utilize a derivatized version of ATP, 6-furfuryl-ATPγS, to transfer thio-phosphates to its substrates (Wan *et al*. 2004). The thiophosphates were then alkylated using *p*-nitrobenzylmesylate to create epitopes that were detected by a thiophosphate ester antibody (Allen *et al*. 2005; Lo and Hollingsworth 2011). 3XFLAG-Rrm3 was purified from vegetative yeast cells as described in (Deegan *et al*. 2019).

GST-Mek1-as autophosphorylation acted as an internal control for the assay (Figure 2A, lane 1). The presence of GST-Mek1-as was confirmed by probing the reaction with α-Mek1 antibodies. Addition of 3XFLAG-Rrm3 resulted in two phosphorylated species (Figure 2A, lane 2). The 3XFLAG-Rrm3 protein was detected using a newly developed antibody against endogenous Rrm3. The specificity of this antibody was confirmed by the absence of a band of the approximate molecular weight for Rrm3 (82 kD) in extracts from a vegetatively grown *rrm3*Δ strain (Figure 2B). Rrm3 phosphorylation was specifically due to GST-Mek1-as, as phosphorylated GST-Mek1-as and 3XFLAG-Rrm3 disappeared when the Mek1-as kinase inhibitor, 1-NA-PP1, was included (Figure 2A, lane 3) (Wan *et al*. 2004). Kinase assays performed in the presence of increasing amounts of bovine serum albumin (BSA) did not exhibit phosphorylated BSA, indicating that GST-Mek1-as does not promiscuously phosphorylate proteins (Figure 2A, lanes 4-6).

**Figure 2.**
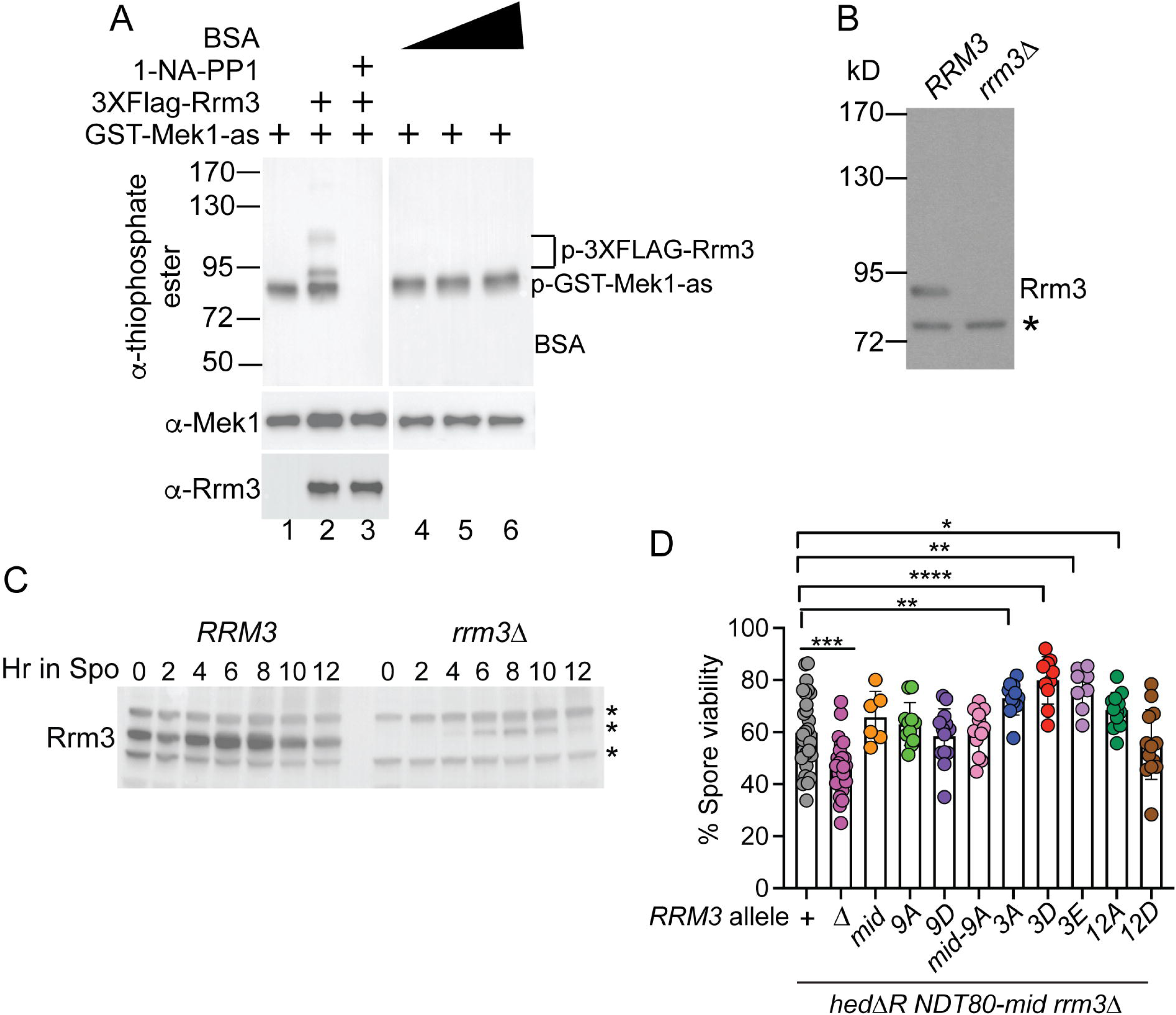
Rrm3 is directly phosphorylated by Gst-Mek1-as in vitro. (A) *In vitro* kinase assays. Kinase assays utilized the semi-synthetic epitope system with 12 nM partially purified GST-Mek1-as (76 kD) and 6-furfuryl-ATPγS. In addition to GST-Mek1-as, 17 nM purified 3XFLAG-Rrm3 (85 kD) was included in the indicated lanes, or increasing amounts (12, 60, 240 nM) of bovine serum albumin (BSA)(66 kD). To inhibit GST-Mek1-as, 15 mM 1-NA-PP1 was used. The top panels were probed with a thio- ester antibody to detect phosphorylated proteins. The presence of GST-Mek1-as and 3XFLAG-Rrm3 in the reactions was confirmed by probing with α-Mek1 and α-Rrm3 antibodies. Numbers on the left are the molecular weights in kiloDaltons (kD) of the prestained markers. The vertical white line indicates the juxtaposition of non-adjacent lanes from the same gel. Horizontal white lines indicate the same samples run on different gels. “p-“ indicates the phosphorylated form of the protein. Numbers at the bottom indicate different lanes. (B) Confirmation of the specificity of the α-Rrm3 antibody. Protein extracts generated from vegetative *RRM3* (NH716) and *rrm3*Δ (NH2485::pRS304^2^) cultures were probed on an immunoblot with α-Rrm3 antibodies. Rrm3 is predicted to have a molecular weight of 82 kD. The asterisk indicates a non-specific band. Numbers indicate the position of the pre-stained molecular weight markers. (C) Rrm3 protein in meiosis. The *RRM3* and *rrm3*Δ diploids from (B) were transferred to Spo medium and samples taken at the indicated timepoints. Protein extracts from each timepoint were probed on immunoblots with α-Rrm3 antibodies. Asterisks indicate non-specific bands detected by the Rrm3 antibody, one of which is induced during meiosis. (D) Complementation of various *rrm3* mutants in the *hed*Δ*R NDT80-mid rrm3*Δ background. The *hed*Δ*R NDT80-mid rrm3*Δ diploid, NH2596, transformed with two copies of *URA3* integrating plasmids containing various alleles of *RRM3*, was sporulated and tetrads dissected to determine the number of viable spores. “Δ” indicates the vector (pRS306), “+” is *RRM3* (pBG22), *RRM3-mid* (pBG22-ΔRPAKR), *RRM3-9A* (pBG22-9A), *RRM3-9D* (pBG22-9D), *RRM3-mid-9A* (pBG22-ΔRPAKR-9A), *rrm3-3A* (pBG22-3A), *rrm3-3D* (pBG22-3D), *rrm3-3E* (pBG22-3E), *rrm3-12A* (pBG22-12A) and *RRM3-12D* (pBG22-12D). Data from the WT diploid (NH716) and *hed*Δ*R NDT80-mid* diploids (NH2505 and NH2610 RCEN) were included as controls. The latter data were previously published in (Ziesel *et al*. 2022). At least 20 tetrads were dissected for each replicate. Dots represent a combination of biological and technical replicates. Statistical significance of differences between strains was determined using the Mann-Whitney test (* = *p*<.05; ** = *p*<.01; *** = *p*<.001, **** = *p*<.0001).

Meiotic timecourses were performed with *RRM3* and *rrm3*Δ diploids to monitor the mobility of the Rrm3 protein during meiosis. Rrm3 protein levels remained fairly constant throughout the wild-type (WT) time course, and no mobility shift indicative of phosphorylation was observed (Figure 2C). Therefore, whether Rrm3 is an *in vivo* substrate of Mek1 remains to be determined.

### The Rrm3-Mek1 interaction is not required for promoting spore viability when Rad51 is constitutively active and the MRC is abolished

To test whether Mek1 phosphorylation of Rrm3 is biologically relevant, various mutant alleles of *RRM3* were tested for complementation of the spore viability defect in the *hed*Δ*R NDT80-mid rrm3*Δ background, where Rad51 is constitutively active and the MRC is inactive (Ziesel *et al*. 2022). *rrm3* alleles containing either a deletion of the RPAKR sequence (*rrm3-mid* for **M**ek1 **I**nteraction **D**efective) or substitutions of the nine N-terminal predicted phosphosites to alanines (*rrm3-9A*) or phosphomimetic aspartates (*rrm3-9D*), fully complemented the *rrm3*Δ, as did the *rrm3-mid-9A* mutant (Figure 2D). Mutants in which the three Mek1 consensus phosphosites in the C-terminal part of Rrm3 were changed to alanines, aspartates or glutamates (another negatively charged amino acid) (*rrm3-3A*, *3D* or *3E*) exhibited a statistically significant increase in spore viability compared to WT (Figure 2D). This increase was also observed in the *rrm3-12A* mutant which has all of the Mek1 consensus phosphosites mutated to alanines, but not the *rrm3-12D* mutant.

### AlphaFold modeling predicts that the Ndt80 RPXKR motif interacts with Mek1 through a negatively charged loop in the FHA domain and an acidic pocket in the kinase domain

Mek1 can be divided into three domains (Niu *et al*. 2007) (Figure 3A). (1) An N-terminal FHA domain. FHA domains canonically mediate protein-protein interactions by binding to phosphorylated threonines, such as the binding of Mek1 to phosphorylated Hop1 in response to DSBs leading to kinase activation (Durocher and Jackson 2002; Carballo *et al*. 2008; Chuang *et al*. 2012). (2) A middle kinase domain. After binding to Hop1, Mek1 activates itself by phosphorylation in *trans* of T327 in the activation loop located within the kinase domain (Niu *et al*. 2007). (3) A C-terminal domain. The function of this part of Mek1 is unknown although indirect genetic evidence suggests that it might promote oligomerization of the kinase (Niu *et al*. 2007).

**Figure 3.**
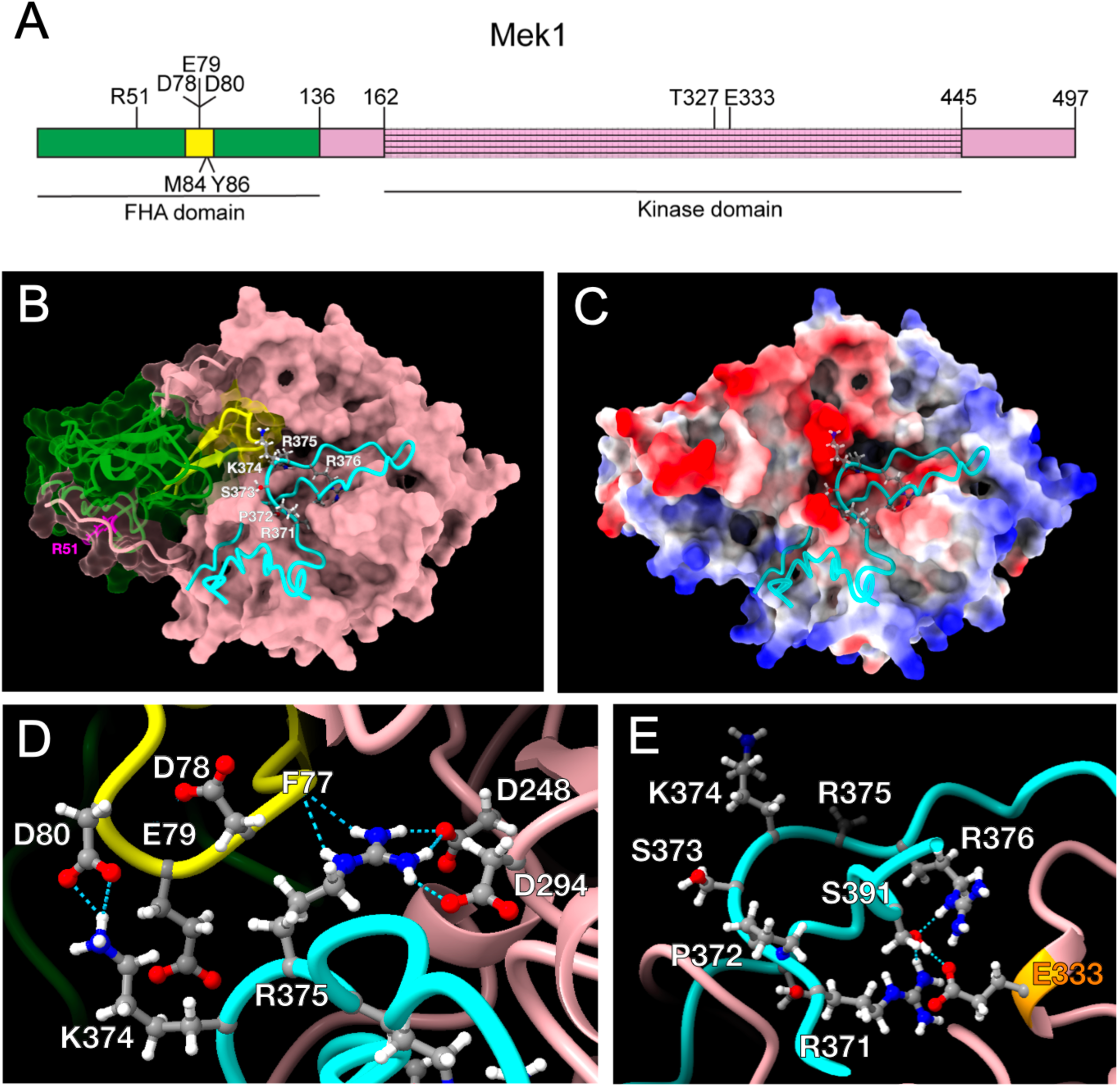
The Mek1 interaction motif in Ndt80 is predicted by AlphaFold Multimer to interact with an acidic sequence within the Mek1 FHA domain and an acidic pocket in the kinase domain. (A) Schematic of the Mek1 protein. The positions of relevant amino acids are indicated. The FHA domain is green while the acidic DED loop flanked by two anti-parallel β-sheets within the FHA domain is colored yellow. The hatched area contains the kinase domain. (B) Structural model generated by AlphaFold Multimer showing the interaction between a 57-amino acid sequence (356-402) from Ndt80 containing the RPSKR Mek1 interaction motif with full-length Mek1. The color code is the same as in A. Cyan indicates the Ndt80 peptide. Magenta represents the arginine at position 51 that required for the FHA domain to bind to phospho-threonine. (C) Surface representation of Mek1 highlighting its electrostatic potential, where red indicates regions of negative charge, blue indicates regions of positive charge, and white represents neutral areas. The Ndt80 peptide is shown in cyan. (D) Close-up of the interaction between Ndt80 and the DED motif of the FHA loop. (E) Close-up of the interaction network between the side chain of Mek1 E333 (orange) and the side chains of Ndt80 R371, R376 and S391.

To determine where the RPSKR motif of Ndt80 interacts with Mek1, AlphaFold structural prediction in multimer mode was used to model the complex between a 57-aa fragment of Ndt80 containing the RPSKR motif and full-length Mek1 (Evans *et al*. 2022). The resulting model indicated that the KR sequence of Ndt80 inserts into an acidic pocket of Mek1, comprising the kinase domain and a negatively charged loop located between two anti-parallel β-sheets in the Mek1 FHA domain (Figure 3BC; Figure S3A). This loop consists of amino acids D78, E79 and D80 (henceforth abbreviated DED).

The model suggests that the side chains of Ndt80 K374 and R375 form ionic interactions with the side chain of D80 and the amide carbonyl oxygen of F77 on the FHA loop. AlphaFold also predicts that Ndt80 R375 makes additional contacts with D248 and D294, contributed by the Mek1 kinase domain. Additionally, E333 in the activation loop of Mek1 forms a network of ionic interactions with Ndt80 R371 (the first arginine in the **R**PSKR motif), R376 (which is immediately downstream of the last arginine of the motif) and S391 (Figure 3E).

Arginine 51 (R51) in Mek1 is analogous to an amino acid in one of the FHA domains of Rad53 that is required for canonical phospho-peptide binding (Durocher *et al*. 1999; Durocher *et al*. 2000). The *mek1-R51A* mutant is defective in kinase activation and exhibits very low spore viability (Niu *et al*. 2007; Chuang *et al*. 2012).

R51 is not, however, required for interaction with Ndt80 (Chen *et al*. 2018). Consistent with this fact, the AlphaFold model places R51 on the other side of the FHA domain from the DED loop, arguing that Mek1 FHA-mediated interaction with Hop1 (which is not a Mek1 substrate) is distinct from that of Ndt80 or Rrm3 (Figure 3B).

AlphaFold Multimer also predicted the structure of Mek1 in complex with a 44-amino acid peptide containing the RPAKR sequence from Rrm3 (Figure S2, S3B). This structure is similar to the Mek1-Ndt80 model, with the KR sequence from Rrm3 located adjacent to the DED acidic loop in the Mek1 FHA domain. In addition, R185 of Rrm3 (equivalent to R371 in Ndt80) is predicted to form hydrogen bonds with E333 in the Mek1 activation loop.

### Mek1 interaction with Ndt80 and Rrm3 requires both the acidic loop in the Mek1 FHA domain and a glutamate in the activation loop

The AlphaFold model suggests that Mek1 interacts with Ndt80 and Rrm3 through a non-canonical FHA domain interface involving the DED loop flanked by two anti-parallel β-sheets. This model is consistent with the fact that mutation of the KR sequence in either *NDT80* or *RRM3* to aspartates and/or alanines disrupts the *lexA-MEK1* interaction (Figure 1C) (Chen *et al*. 2018). This hypothesis is further supported by the observation that mutation of conserved amino acids located on the flanking anti-parallel β-sheets to alanine, (*mek1-M84 Y86* and *mek1-W73 V75*) disrupted the two-hybrid interaction with *GAD-NDT80^284-627^*without affecting Mek1 steady state protein levels (Figure S4BC).

The AlphaFold model predicts that mutation of the DED sequence should specifically disrupt *lexA-MEK1* interaction with *GAD-NDT80^284-627^* and *GAD-RRM3^51-723^* (hereafter *GAD-NDT80* and *GAD-RRM3*, respectively), without affecting the two-hybrid signal for *GAD* fusions such as *GAD-SRS2^1035-1174-^*^Δ*RKSKR*^, *GAD-SGS1^82-614^*and *GAD-MMS4^91-378^* that lack an RPXKR motif. Single amino acids substitutions within the DED sequence to alanine (*D78A*, *E79A*, or *D80A*) had no obvious effect on interactions with any of the *GAD* fusion proteins (Figure S1AB). In contrast, substitution of the DED amino acids to alanine in *lexA-mek1-D78A E79A D80A* (hereafter *lexA-mek1-AAA*) specifically abolished the two-hybrid signal with *GAD-NDT80* and *GAD-RRM3* (Figure 4A). Steady state lexA-Mek1-AAA protein levels were not reduced compared to lexA-Mek1, ruling out protein instability as a trivial explanation for the defect in Ndt80 and Rrm3 interaction (Figure 4B).

**Figure 4.**
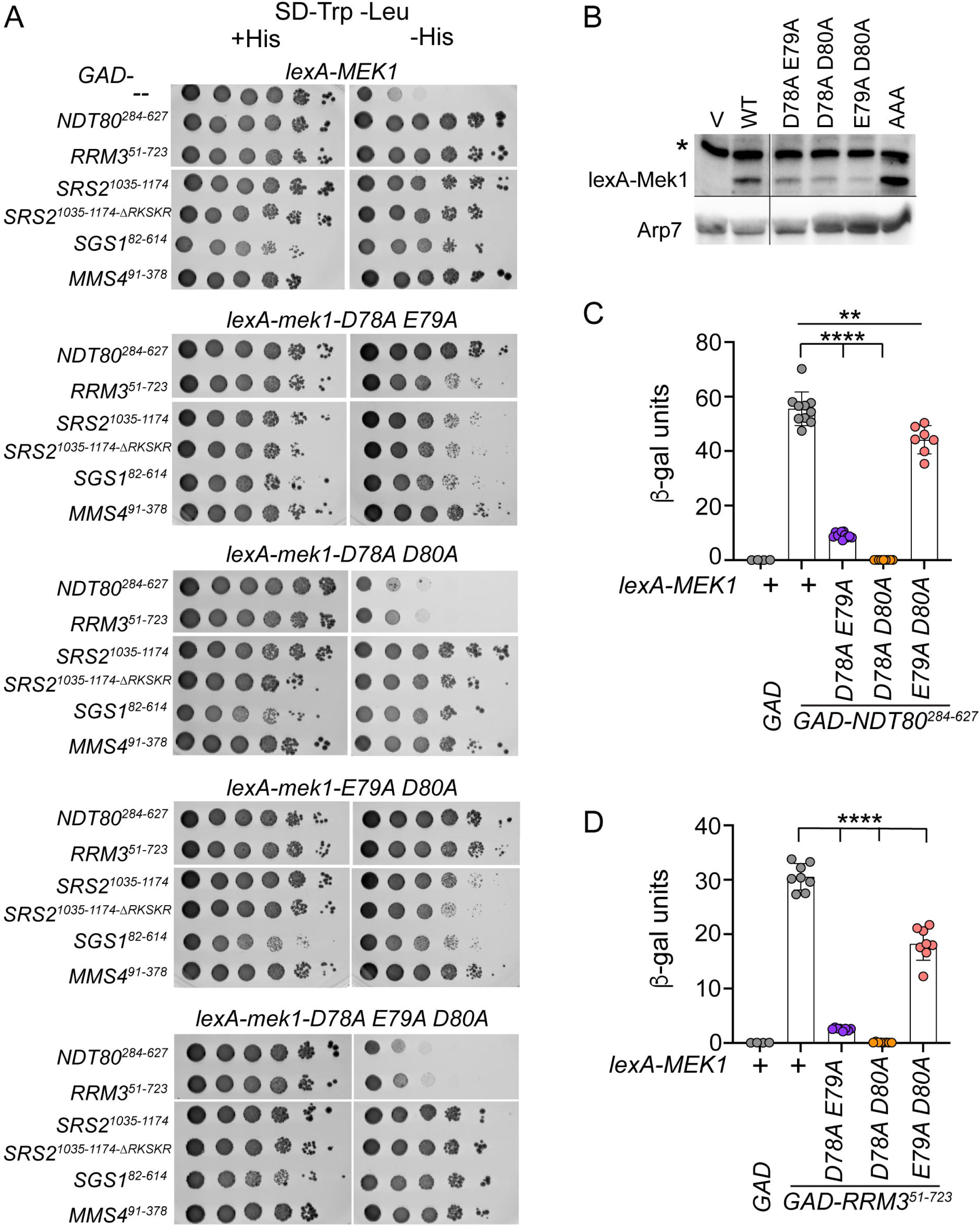
An acidic patch in the Mek1 FHA domain is required specifically for interaction the two-hybrid interaction between Ndt80 and Rrm3 in the two-hybrid system. (A) Two-hybrid interactions detected by spotting assays. A *lexA-MEK1* plasmid (pTS3) or its indicated mutant derivatives was co-transformed with *LEU2* plasmids containing either *GAD* (pACTII), *GAD-NDT80^284-627^* (pXC13), *GAD-RRM3^51-723^* (pJW1), *GAD-SRS2^1035-1174^*(pGAD-SRS2), *GAD-SRS2^1035-1174^* ^Δ*RKSKR*^ (pGAD-SRS2-ΔRKSKR), *GAD-SGS1^82-614^* (pA46), or *GAD-MMS^491-378^* (pA8) into L40. Spotting assays were performed as described in Figure 1. (B) Immunoblots of protein extracts generated from L40 transformed with a plasmid carrying the indicated *lexA-MEK1* allele were probed with α-Mek1 or α-Arp7 antibodies. “V” indicates the *lexA* plasmid, pSTT91. The vertical black line indicates where the blots were cut and an irrelevant lane was omitted. The horizontal black line indicates the juxtaposition of the Arp7 blot with the lexA-Mek1 blot. The asterisk indicates a non-specific band. (C) Two-hybrid interactions detected by liquid β-galactosidase assays using L40 cultures containing the indicated *lexA-MEK1* alleles (pTS3-D78A E79A, pTS3-D78A D80A, or E79A D80A) combined with *GAD-NDT80^284-627^*. Each dot represents an independent transformant for which the average value from two technical replicates was calculated. “+” indicates WT. (D) Same as Panel C except using *GAD-RRM3^51-723^*. Statistical significance was determined using an unpaired Student’s t-test: ** = *p* = 0.0012 and **** = *p* < 0.0001. The *lexA-MEK1/GAD* data are the same in both graphs.

Double mutant combinations of the DED sequence varied in their *lexA-MEK1* interaction phenotypes (Figure 4A). While *lexA-mek1-D78A D80A* resembled *lexA-mek1-AAA*, *lexA-mek1-D78A E79A* and *lexA-mek1-E79A D80A* appeared to interact similarly to *lexA-MEK1* when expression of the *HIS3* reporter was assayed by growth on medium lacking histidine. However, because phenotypic assays described below suggested that there was some impairment in the binding of these *lexA-mek1* mutants to *GAD-NDT80,* expression of the *lacZ* reporter gene was monitored using liquid β-galactosidase assays as these are more sensitive and quantitative. These experiments verified that the *lexA-mek1-D78A E79A* and *E79A D80A* double mutants exhibit weaker two-hybrid interactions with *GAD-NDT80* and *GAD-RRM3* compared to *lexA-MEK1* (Figure 4CD).

A second prediction of the AlphaFold structures is that E333 in the Mek1 activation loop interacts with the Ndt80 and Rrm3 RPXKR sequences. This prediction was supported by showing that *lexA-mek1-E333A* and *lexA-mek1-E333K* disrupted the two hybrid interactions with *GAD-NDT80* and *GAD-RRM3*, without affecting binding to the other *GAD* fusions or steady state proteins levels (Figure S1CD). Together, these analyses validate the AlphaFold models, and suggest that, in addition to its canonical function in binding to phosphopeptides, the Mek1 FHA domain non-canonically interacts with a subset of Mek1 substrates through an RPXKR motif.

### The Mek1 FHA domain DED sequence is not required for kinase activation

Because binding of the Mek1 FHA domain to phosphorylated Hop1 is required for kinase activation, it is possible that mutations in this domain disrupt the structure of the FHA domain. If true, not only would the Ndt80/Rrm3 interaction with Mek1 be lost, but so would the ability of Mek1 to bind to phospho-Hop1. In this case, it would not be possible to distinguish whether meiotic phenotypes were due to failure of Mek1 to bind to a particular protein or because the kinase was inactive. Mek1 kinase activation was assessed by detection of phosphorylated T327 in the Mek1 activation loop (Niu *et al*. 2007; Wu *et al*. 2010). These experiments were performed in an *ndt80*Δ diploid that causes cells to arrest in pachytene with fully synapsed chromosomes (Xu *et al*. 1995). Although chromosome synapsis results in the removal of the bulk of Mek1 and the recombination machinery, a low number of DSBs continue to be made, resulting in a low level of Mek1 kinase activity (Subramanian *et al*. 2016; Prugar *et al*. 2017; Mu *et al*. 2020). *mek1*Δ diploids progress faster through meiosis than *MEK1* and the advantage of the *ndt80*Δ diploid is that cells can accumulate at the arrest point, making it easier to compare different *mek1* mutants (Malone *et al*. 2004). Phosphorylated T327 was observed in the *mek1* DED double and triple mutant combinations, indicating that this sequence is not required for kinase activation (Figure 5A). In contrast, the signal for the Mek1-R51A protein was reduced compared to WT, as was the steady state level of the protein (Figure 5A). Amino acid substitutions in one of the flanking β-sheets (*mek1-M84A Y86A*) also exhibited a low level of T327 phosphorylation. One possibility is that the β-sheet mutations create a conformational change within the FHA domain that prevents both interaction with Ndt80 and binding to Hop1.

**Figure 5.**
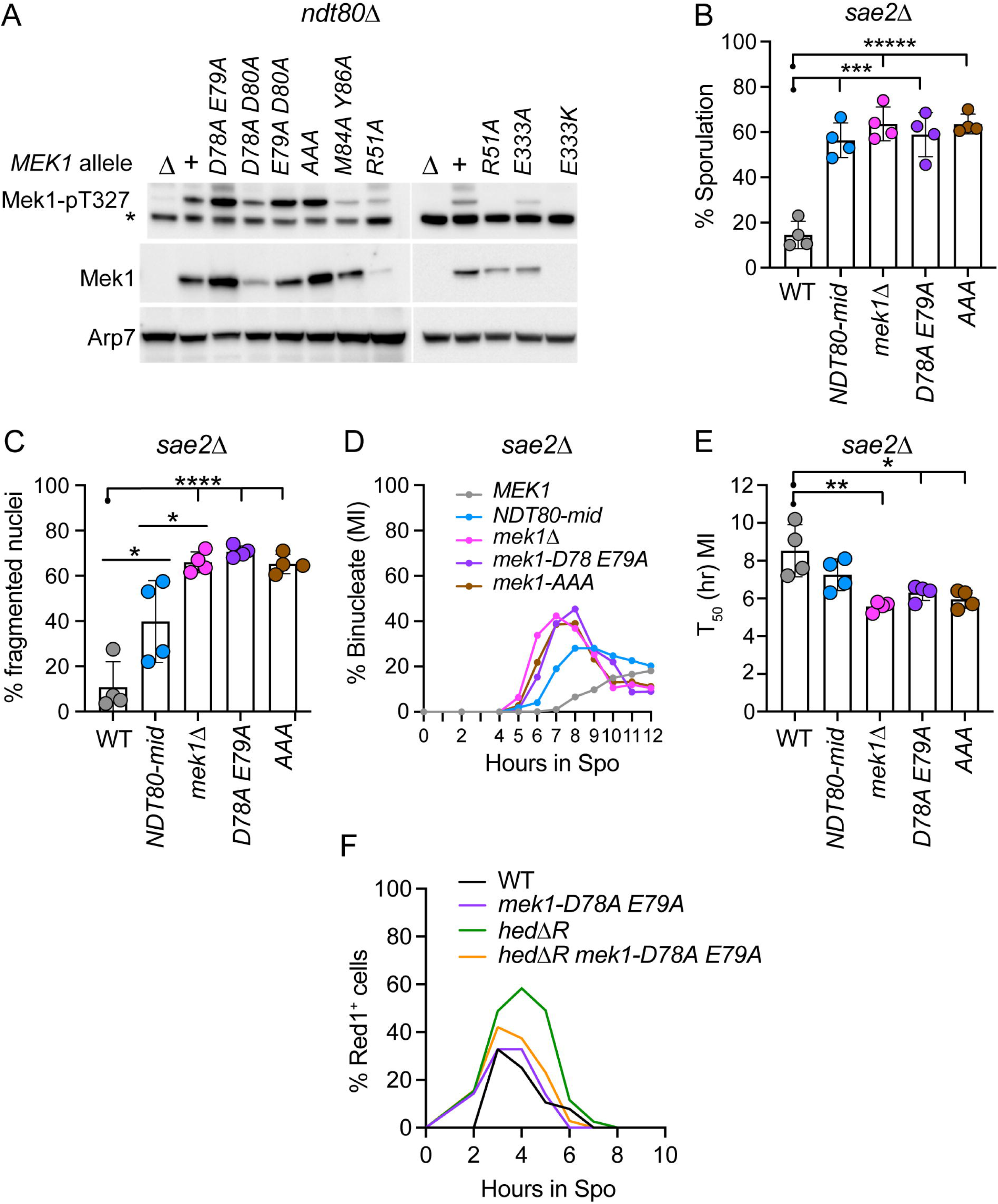
The Mek1 FHA domain DED sequence is not required for kinase activation but is necessary for the meiotic recombination checkpoint. (A) *mek1*Δ *ndt80*Δ diploids (NH2750) homozygous for either of the following *mek1* alleles, *mek1*Δ (pRS306), *MEK1* (pLP37), *mek1-D78A E79A* (pLP37-D78A E79A), *mek1-D78A D80A* (pLP37-D78A D80A), *mek1-E79A D80A* (pLP37-E79A D80A), *mek1-AAA* (pLP37-D78A E79A D80A), *mek1-M84A Y86A* (pLP37-MY), or *mek1-R51A* (pLP37-R51A) were incubated in Spo medium for six hours. Mek1 activity was monitored by detection of phosphorylated T327 using an antibody against phospho-Akt substrates (Niu *et al*. 2007; Wu *et al*. 2010). Total Mek1 and Arp7 were used as loading controls. (B) MRC function: sporulation. *mek1*Δ *sae2*Δ diploids (NH2758) homozygous for either *mek1*Δ, *MEK1*, *mek1-D78A E79A*, or *mek1-AAA* were transferred to liquid Spo medium and incubated at 30°C for >24 hours. For each replicate, 200 cells were examined by light microscopy for the presence of asci. The *NDT80-mid sae2*Δ diploid (NH2759::pNH317^2^) was included for comparison. Dots indicate different biological replicates (*n* = 4). Error bars indicate the means and standard deviations. A Black dot indicates the strain to which the other strains were compared. (C) MRC function: nuclear fragmentation. Cells from the cultures shown in panel B were taken at the 12-hour timepoint, fixed and stained with DAPI and the number of cells exhibiting >4 DAPI staining foci out of 200 were counted. (D) MRC Function: Progression through Meiosis I. DAPI stained cells from the indicated timepoints were counted for the number of binucleate cells out of 200. Lines represent the average % binucleate cells for each timepoint (*n* = 4). (E) T_50_ analysis. T_50_ indicates the time in hours that it took the sporulating cultures from Panel D to reach half maximal % binucleate value. Data were analyzed for statistical significance using an unpaired, two-tailed Student’s t test. (F) Red1 immunofluorescence. *MEK1* (NH729::pLP37^2^), *mek1-D78A E79A* (NH729::pLP37-D78A E79A^2^), *hed*Δ*R* (NH2603::pLP37^2^) and *hed*Δ*R mek1-D78A E79A* (NH2603::pLP37-D78A E79A^2^) diploids were sporulated, cells fixed at the indicated timepoints and stained with Red1 antibodies. The graph shows the average values at each time point for two biological replicates. For all graphs, * = *p* < 0.05; ** = *p* < 0.01, **** = *p*< 0.0001.

Mek1 E333 is required for interaction with both Ndt80 and Rrm3 (Figure S1C). This amino acid is close to T327 and T331, both of which are phosphorylated *in vivo* and are located within the Mek1 activation loop (Figure 3A) (Niu *et al*. 2007). In meiotic yeast cells, the Mek1-E333K protein was unstable, while the Mek1-E333A protein accumulated to a level similar to Mek1-R51A (Figure 5A). Only a low level of Mek1 T327 phosphorylation was observed in the *mek1-E333A* diploid (Figure 5A). Therefore like the *mek1-MY* mutant, phenotypes observed for *mek1-E333A* are most likely due to a disruption of kinase activation as opposed to a defect in a specific protein-protein interaction.

### The Mek1 FHA domain DED sequence is required for the meiotic recombination checkpoint

Disrupting the interaction between Mek1 and Ndt80 by deleting the Ndt80 RPSKR sequence impairs the MRC (Chen *et al*. 2018). If the failure of the *lexA*-*mek1-AAA* mutant to interact with *GAD-NDT80* in the two-hybrid system reflects what is happening during meiosis (i.e, the Mek1-AAA protein is unable to bind to Ndt80), the *mek1-AAA* mutant should also be defective in the MRC. MRC activity can be measured using mutants such as *rad50S* or *sae2*Δ*/com1*Δ in which DSBs are made but not resected (Alani *et al*. 1990; Mckee and Kleckner 1997; Prinz *et al*. 1997). The unresected DSBs cannot be repaired and therefore persist, keeping Mek1 active for longer. As a result, meiotic progression is delayed and sporulation is reduced in a *MEK1*-dependent manner (Xu *et al*. 1997).

Inactivation of the MRC, either by *NDT80-mid* or *mek1*Δ, suppressed the *sae2*Δ sporulation defect, increasing from ∼15% to ∼60% asci (Figure 5B). The *mek1-AAA* mutant exhibited this suppression as well, indicating it is also defective in the MRC (Figure 5B). Interestingly, *mek1-D78A E79A* suppressed the *sae2*Δ sporulation defect to the same extent as *mek1-AAA*, even though it exhibited a stronger Ndt80 interaction signal in the two-hybrid assay (Figure 4A, 5B). Meiotic progression can be followed using the DNA-staining fluorescent molecule, 4’.6-diamidino-2-phenylindole (DAPI). It was previously observed that at later timepoints in an *sae2*Δ diploid some of the nuclei “degenerated” as a result of chromosome fragmentation (Mckee and Kleckner 1997). This fragmentation (i.e., nuclei with > four DAPI foci) was exacerbated when the MRC was compromised, as expected if cells progressed through meiosis with unrepaired DSBs (Figure 5C). *mek1-AAA* and *mek1-D78A E79A* exhibited similar numbers of fragmented nuclei as *mek1*Δ, providing additional support that they are defective in the MRC. The *NDT80-mid sae2*Δ did not produce as many fragmented nuclei as the *mek1* mutants (Figure 5C). This could be because *NDT80-mid* does not inactivate the MRC as efficiently as eliminating *MEK1*. Alternatively, the *mek1*Δ, *mek1-AAA* and *mek1-D78A E79A* mutants could be pleiotropic and affect other processes that still occur in the *NDT80-mid* strain which delay exit from prophase I.

Mek1 inhibition of Ndt80 prevents exit from pachytene by preventing expression of Ndt80 targets such as *CDC5* and *CLB1*, which are important for resolving double Holliday junctions, disassembling synaptonemal complexes and building the Meiosis I spindle (Sourirajan and Lichten 2008; Okaz *et al*. 2012; Chen *et al*. 2018). Therefore *sae2*Δ cells defective in the MRC should undergo Meiosis I sooner than when the MRC is functional. In fact, *mek1*Δ *sae2*Δ, *mek1-AAA sae2*Δ and *mek1-D78A E79A* all completed Meiosis I significantly faster than *sae2*Δ alone (Figure 5DE). The T_50_ value measures the length of time it takes a culture to reach the half maximal number of binucleate cells. This number was underestimated for the *sae2*Δ diploid because the number of binucleate cells had not peaked by 12 hours (Figure 5D). Meiosis I occurred earlier in the *NDT80-mid sae2*Δ as well, although not as early as the *mek1* mutants (Figure 5D). The difference between *NDT80-mid sae2*Δ and *sae2*Δ was not statistically significant, which may be due to the underestimation of the *sae2*Δ T_50_ value (Figure 5E).

Recently, it was shown that constitutively activating Rad51 in the *hed*Δ*R* background triggers an MRC-dependent delay in exiting from prophase I (Ziesel *et al*. 2022). Red1 is a meiosis-specific axial element protein that is degraded when Ndt80 is activated (Rockmill and Roeder 1988; Thompson and Roeder 1989; Smith and Roeder 1997; Sourirajan and Lichten 2008). Prophase exit can therefore be monitored using whole cell immunofluorescence that detects Red1 (Ziesel *et al*. 2022). The frequency of Red1 positive cells was determined at different times in meiotic timecourses comparing WT, *mek1-D78A E79A*, *hed*Δ*R*, and *hed*Δ*R mek1-D78A E79A*. Consistent with previous results, it took longer for the percentage of Red1^+^ cells to decrease in the *hed*Δ*R* diploid compared to the other three strains (Figure 5F) (Ziesel *et al*. 2022). In contrast, although *hed*Δ*R mek1-D78A E79A* entered prophase I with similar kinetics as *hed*Δ*R*, it exited prophase I sooner, resembling instead the WT and *mek1-D78A E79A* diploids (Figure 5F). The fact that *mek1-D78A E79A* is defective in the MRC when triggered by two different mutants supports the hypothesis that these mutations are disrupting interaction with Ndt80 as predicted by the AlphaFold model.

### The Mek1 FHA acidic loop is required to promote meiotic chromosome segregation and spore viability

One of Mek1’s critical functions is to bias strand invasion into homologs instead of sister chromatids. In the absence of *MEK1* activity, this bias is reversed and the lack of interhomolog crossovers causes high levels of aneuploid inviable spores (Rockmill and Roeder 1991; Leem and Ogawa 1992; Niu *et al*. 2005; Kim *et al*. 2010). Various *mek1* mutants were introduced into an otherwise wild-type strain and tetrads dissected to determine the frequency of viable spores. Mutants that exhibited decreased kinase activation (*mek1-M84A Y86A, R51A* and *E333A*) exhibited a corresponding decrease in spore viability (Figure 5A, 6A). Of the DED single and double mutants, only *mek1-D78A D80A* had statistically significant lower spore viability compared to WT (Figure 6A). This phenotype could be due to the lower steady state protein levels exhibited by the Mek1-D78A D80A protein in the *ndt80*Δ background (Figure 5A). The *mek1-AAA* had greatly reduced spore viability compared to WT (∼20% compared to ∼95%) (Figure 6A). This phenotype cannot simply be explained by its defects in binding to Ndt80 or Rrm3, as the *NDT80-mid*, *rrm3*Δ and *NDT80-mid rrm3*Δ mutants displayed nearly WT spore viability (Figure 6C) (Ziesel *et al*. 2022).

**Figure 6.**
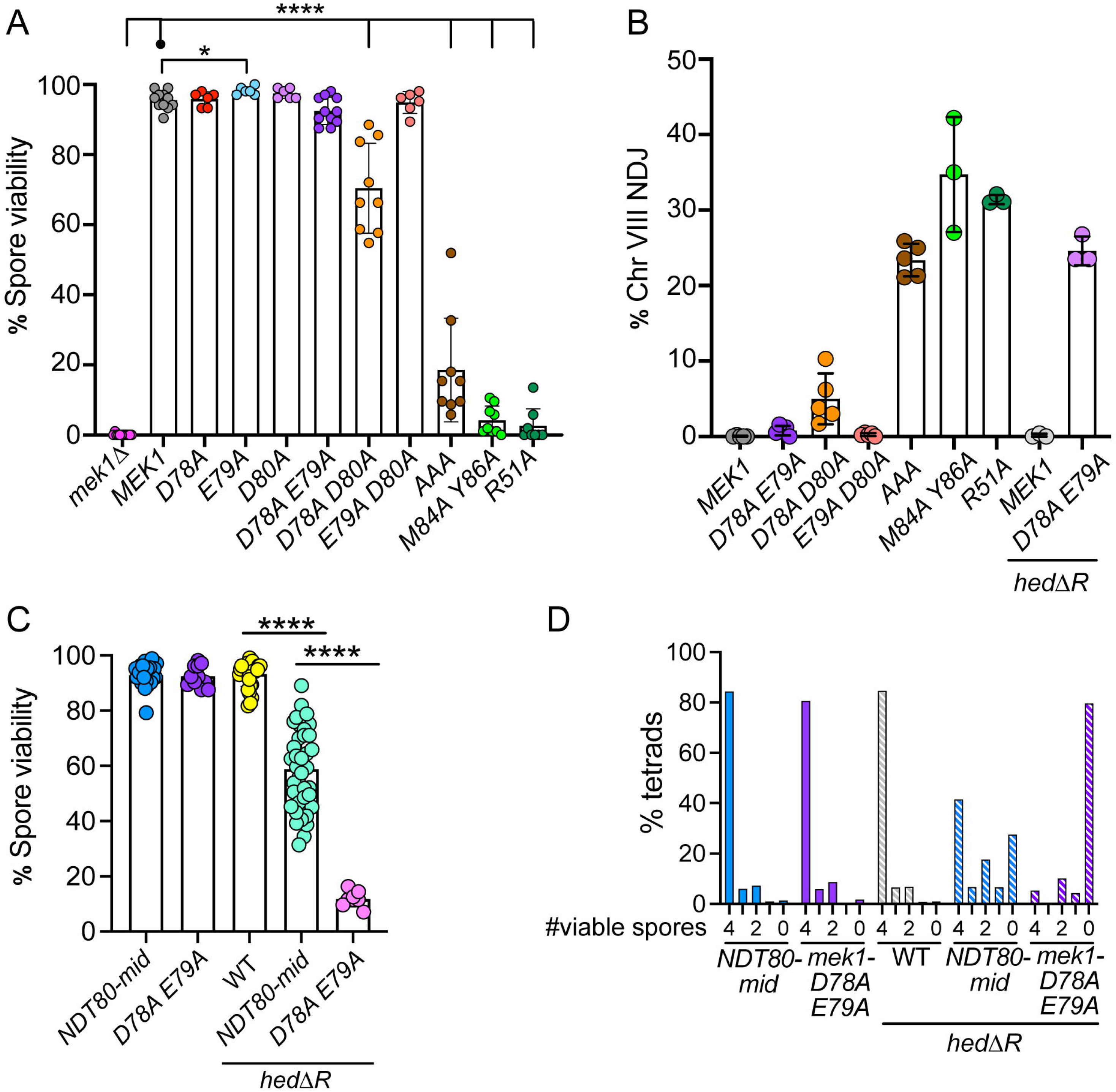
Phenotypic characterization of *mek1* FHA domain mutants in meiosis. (A) Spore viability. Isogenic *mek1*Δ diploids, (NH2740 RCEN, NH2740 RC20, NH729) homozygous for a plasmid containing *MEK1* (pLP37), or the indicated mutant *mek1* alleles, were sporulated and tetrads dissected to determine spore viability. The line with a dot indicates the strain to which the other strains were compared. Only statistically significant comparisons are shown. The analysis was done as described in Figure 2D. (B) Chromosome VIII non-disjunction. The NH2740 RCEN diploids used in Panel A contain *RFP* and *CFP* integrated at allelic positions adjacent to the centromeres of chromosome VIII (Thacker *et al*. 2011). These diploids were sporulated in liquid medium and the tetrads analyzed by fluorescence microscopy for chromosome VIII segregation. Each dot indicates a biological replicate. The total number of tetrads scored was: *MEK1* (1879), *mek1-D78A E79A* (1457), *mek1-D78A D80A* (1743), *mek1-E79A D80A* (2005), *mek1-AAA* (914), *mek1-M84A Y86A* (466), and *mek1-R51A* (304). The latter two mutants behaved like *mek1*Δ and the spores were difficult to score for fluorescence. The *hed*Δ*R mek1*Δ diploids homozygous for either *MEK1* or *mek1-D78A E79A* (NH2783 RCEN::pLP37^2^ and NH2783 RCEN::pLP37-D78A E79A^2^, respectively) were similarly scored (683 tetrads for *MEK1*, 444 tetrads for *mek1-D78A E79A*). Error bars show the means and standard deviations. (C) Spore viability in the *hed*Δ*R* background. The *mek1-D78A E79A* (NH2740::pLP37^2^), *hed*Δ*R* (NH2603::pLP37^2^) and *hed*Δ*R mek1-D78A E79A* (NH2603::pLP37-D78A E79A^2^) diploids were sporulated and tetrads dissected. Spore viability data from the *NDT80-mid* and *hed*Δ*R NDT80-mid* diploids were previously published in (Ziesel *et al*. 2022). The *mek1-D78A E79A* data are repeated from Panel A for comparison. (D) Distribution of viable spores in tetrads. The percent tetrads with 4, 3, 2, 0 or 1 viable spores was plotted using the data from Panel C. Meiosis I non-disjunction is indicated by a decrease in 4 viable:0 inviable tetrads and an increase in 2:2 and 0:4 tetrads. Statistical analyses for the graphs in this figure used the Mann-Whitney test. (* = *p* < 0.05; ** = *p* < 0.01; *** = *p* < 0.001, **** = *p* < 0.0001).

If the spore inviability observed for *mek1-AAA* was due to a loss of interhomolog bias, then the resulting lack of interhomolog crossovers due to increased intersister recombination should result in increased levels of Meiosis I non-disjunction. Segregation of chromosome VIII was therefore monitored using a fluorescent spore assay (Thacker *et al*. 2011). Genes encoding a red fluorescent protein (*RFP*) or a cyan fluorescent protein (*CFP*) under control of a spore autonomous promoter were integrated at allelic positions adjacent to the centromeres of chromosome VIII. Proper segregation produced tetrads with two red and two blue spores, while tetrads in which Meiosis I non-disjunction of chromosome VIII occurred exhibited two pink and two black spores (Thacker *et al*. 2011; Ziesel *et al*. 2022). The benefits of this system are that tetrad dissection is not required and spores do not need to be viable to be scored. The severity of the spore viability defect for each mutant was mirrored by an increase in Meiosis I non-disjunction. For example, *mek1-D78A E79A* and *E79A D80A* had nearly WT spore viability and very low levels of Meiosis I non-disjunction, while chromosome VIII mis-segregated in ∼30% of the asci from the *mek1-M84A Y86A* and *R51A* diploids (Figure 6B). Meiosis I non-disjunction was also significantly increased in *mek1-AAA*, suggesting that disrupting the ability of Mek1 to interact with proteins containing an RPXKR motif interferes with proper chromosome segregation at the first meiotic division.

### The Mek1 FHA domain acidic loop promotes crossover formation when Rad51 is constitutively active during meiosis

The phenotypes of the *mek1-D78A E79A* mutant are similar to *NDT80-mid*: *i.e*., nearly WT spore viability is coupled with a defect in the MRC (Figure 5, 6A) (Chen *et al*. 2018). When *RAD51* is constitutively active during meiotic prophase in the *hed*Δ*R* background, formation of interhomolog crossovers formed by Rad51-mediated strand invasion requires time provided by the MRC, as evidenced by a decrease in spore viability in a *hed*Δ*R NDT80-mid* diploid (Figure 6C) (Ziesel *et al*. 2022). If *mek1-D78A E79A* is only defective in Ndt80 interaction, then combining it with *hed*Δ*R* should phenocopy the *hed*Δ*R NDT80-mid* diploid. This was not the case, however. Spore viability in the *hed*Δ*R mek1-D78A E79A* diploid was reduced to ∼15% compared to the ∼60% average spore viability of *hed*Δ*R NDT80-mid* (Figure 6C). The *hed*Δ*R mek1-D78A E79A* spore inviability was due to Meiosis I non-disjunction, as evidenced by a dramatic increase in Chromosome VIII mis-segregation and that the distribution of viable spores in tetrads exhibited a Meiosis I non-disjunction pattern (Figure 6BD) (Hollingsworth *et al*. 1995).

A defect in interhomolog bias results in Meiosis I nondisjunction because interhomolog crossovers are necessary to connect the homologs. Both crossover (CO) and non-crossover (NCO) interhomolog recombination was therefore analyzed in WT, *mek1-D78A E79A*, *hed*Δ*R* and *hed*Δ*R mek1-D78A E79A* diploids using physical analysis of the *HIS4LEU2* hotspot (Hunter and Kleckner 2001; Wu *et al*. 2010). The DSB site in this hotspot is flanked by asymmetrically located XhoI sites on the two homologs, with an NgoMIV restriction site located adjacent to the DSB site on one of the homologs (Figure 7A). Digestion of the genomic DNA with both NgoMIV and XhoI produced diagnostic fragments for a subset of crossovers (CO2) and noncrossovers (NCO1). Consistent with the literature a ∼2 fold reduction in both CO and NCOs was observed in the *hed*Δ*R* diploid (Figure 7BCD) (Liu *et al*. 2014; Ziesel *et al*. 2022). The *mek1-D78A E79A* diploid also exhibited a ∼2 fold reduction in COs, indicating that *MEK1* function is affected but not to the extent that spore viability is reduced. A greater reduction in both NCOs and COs was observed in the *hed*Δ*R mek1-D78A E79* diploid (Figure 7BCD). The interhomolog recombination defect of the *hed*Δ*R mek1-D78A E79* mutant suggests that Rad51-mediated interhomolog recombination is more sensitive to the amount of Mek1 function than Dmc1-mediated recombination.

**Figure 7.**
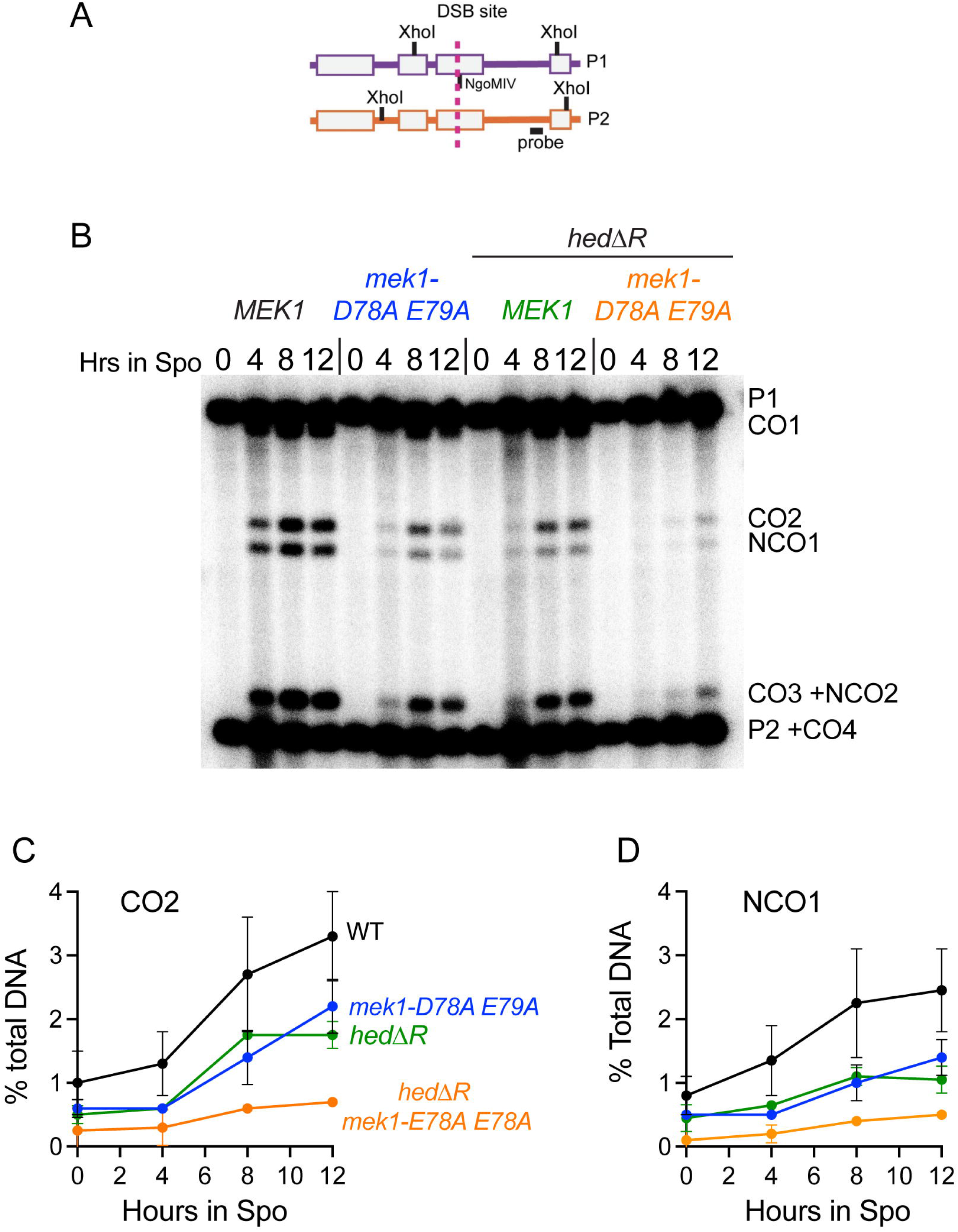
Interhomolog recombination at the *HIS4LEU2* hotspot in *mek1-D78A E79A* and *hed*Δ*R mek1-D78A E79A* diploids. (A) Schematic of the *HIS4LEU2* hotspot (Ziesel *et al*. 2022). The vertical dotted line indicates where Spo11-mediated DSBs occur. The black box indicates the position of the probe. (B) Southern blot. Genomic DNA from WT (NH729::pLP37), *mek1-D78A E79A* (NH729::pLP37-D78A E79A^2^), *hed*Δ*R* (NH2603::pLP37) and *hed1*Δ*R mek1-E78A E79A* (NH2603::pLP37-D78A E79A) at the indicated timepoints was digested with XhoI and NgoMIV. P1/P2 indicate parental bands, CO1/CO2/CO3/CO4 indicate different crossover bands, and NCO1/NCO2 indicate different noncrossover bands. (C) Quantification of the CO2 band. (D) Quantification of the NCO1 band.

## Discussion

### The Rrm3 helicase is an in vitro substrate of Mek1

The Rrm3 helicase promotes interhomolog recombination when Rad51 is constitutively activated during meiosis (Ziesel *et al*. 2022). Rad51 strand exchange activity is normally inhibited by Mek1 while interhomolog recombination is occurring and so a reasonable hypothesis is that Rrm3 is similarly inhibited by Mek1 (Niu *et al*. 2009; Callender *et al*. 2016). The Rrm3 helicase resembles Ndt80 in that it is an *in vitro* substrate of Mek1 and it interacts with the Mek1 FHA acidic loop through an RPXKR sequence. In vegetative cells, the Mek1 paralog Rad53, phosphorylates and inhibits Rrm3 in response to replication stress (Rossi *et al*. 2015). While the Rrm3 N-terminal domain is not required for helicase activity *in vitro* (Ivessa *et al*. 2002), it does play a role in regulating Rrm3 stability, replication fork progression through discrete sites, including G4 quadruplex structures, as well as viability when DNA repair is compromised (Ivessa *et al*. 2003; Bessler and Zakian 2004; Varon *et al*. 2024). The observation that this domain contains multiple Mek1 consensus phosphorylation sites and the RPAKR Mek1 interaction sequence makes it an intriguing target for regulation by Mek1. However, no obvious phenotypes were observed even when the RPAKR sequence was deleted in combination with alanine or aspartate mutations in the putative N-terminal Mek1 phosphosites. In contrast, mutation of the three C-terminal Mek1 consensus sites significantly increased spore viability in the *hed*Δ*R* background. One possibility is that Mek1 phosphorylation of the putative C terminal phosphosites inhibits Rrm3, but the D and E mutants do not function as good phosphomimics. It is also possible that the increase in spore viability has nothing to do with phosphorylation, but instead Rrm3 function is improved for unknown reasons by changing the amino acids located at the C terminus. Therefore whether Mek1 phosphorylation of Rrm3 has a biological function during meiosis is not yet clear.

### The Mek1 FHA domain binds non-canonically to RPXKR motifs in a subset of its substrates

Although consensus sites surrounding phosphorylation sites have been determined for many kinases, this information is usually not sufficient to provide the specificity needed for a kinase to phosphorylate specific targets (Mok *et al*. 2010; Miller and Turk 2018). Kinase specificity can be enhanced by binding of peptide motifs on a substrate either within the catalytic domain or in a separate domain of the kinase (Miller and Turk 2018). A well-studied example of the latter situation is the polo-box domain (PBD) found in the family of polo-like kinases. PBDs are usually located in the C-terminal part of the protein, separate from the kinase domain, and they bind to phosphorylated motifs on target proteins (Park *et al*. 2010). Recent work has shown that PBDs can also mediate protein-protein interactions that are independent of phosphorylation. For example, a region on the PBD from the yeast polo-like kinase, Cdc5, located opposite the phospho-peptide recognition motif, binds non-canonically to conserved sequences on Dbf4 and Exo1 (Chen and Weinreich 2010; Almawi *et al*. 2020; Sanchez *et al*. 2020).

FHA domains can also mediate protein-protein interactions by binding to phosphorylated threonines on target proteins, but unlike PBDs, FHA domains are not restricted to protein kinases (Durocher *et al*. 2000; Almawi *et al*. 2017). The Mek1 FHA domain serves a canonical purpose in the activation of Mek1 kinase activity by binding to phospho-threonines on Hop1, thereby localizing Mek1 to the axis near DSBs (Carballo *et al*. 2008; Chuang *et al*. 2012; Kniewel *et al*. 2017). Mek1 autophosphorylation in *trans* of T327 in the activation loop then activates the kinase (Niu *et al*. 2007). Phosphorylation-independent binding between FHA domains and other proteins has previously been observed (Almawi *et al*. 2017). For example, the Mek1 paralog, Rad53, interacts with Dbf4 through a non-canonical interface on one of the Rad53 FHA domains that does not involve the phosphate moiety (Matthews *et al*. 2014).

This work reveals a different non-canonical way in which FHA domains can mediate protein-protein interactions. AlphaFold Multimer predicted that the positively charged KR amino acids in the conserved RPxKR motifs of Ndt80 and Rrm3 are in the vicinity of an acidic loop (D78 E79 D80) within the Mek1 FHA domain. This model was validated by showing that mutation of these amino acids to alanine specifically disrupted two-hybrid interactions with *GAD-NDT80* and *GAD-RRM3*, but not with *GAD* fusions that do not contain an RPxKR motif. The acidic loop is located on the opposite side of the Mek1 FHA domain relative to arginine 51, which is required for binding to phosphorylated Hop1 but not to Ndt80 (Niu *et al*. 2007; Chuang *et al*. 2012; Chen *et al*. 2018). Importantly, mutations in the acidic loop do not affect kinase activation, as indicated by phosphorylation of T327 in the Mek1 activation loop. Therefore mutant phenotypes can be interpreted as being caused by defective protein-protein interactions, rather than by alterations in the FHA domain structure that prevent binding to phospho-Hop1.

AlphaFold also predicted an interaction between the RPxKR sequences of Ndt80 and Rrm3 with E333 in the Mek1 activation loop. The *lexA-mek1-E333A* and *lexA-mek1-E333K* mutants both specifically abolished the two-hybrid signal with *GAD-NDT80* and *GAD-RRM3*, validating the model. However, these mutants also affected kinase activation, perhaps due to their proximity to T327 and T331, both of which promote kinase activation and so were not pursued further (Niu *et al*. 2007).

### The acidic loop in the Mek1 FHA domain is required for the meiotic recombination checkpoint

We have been unable to directly test the effects of the *NDT80-mid* and *mek1* DED mutants on Mek1-Ndt80 complex formation in meiotic cells due to insolubility of the Ndt80 protein (unpublished observations). Instead, we have indirectly tested the AlphaFold model by phenotypic characterization of the *mek1-D78A E79A* and *mek1-AAA* mutants with regard to the MRC.

The *NDT80-mid* allele disrupts the Mek1-Ndt80 interaction because it lacks the RPSKR sequence on the transcription factor (Chen *et al*. 2018). As a result, Ndt80 is not phosphorylated by Mek1 and the MRC is no longer functional (Chen *et al*. 2018). If mutations in the Mek1 acidic loop affect binding to Ndt80 in meiotic cells, a similar phenotype should be observed. This was, indeed, the case. MRC activity was monitored using the *sae2*Δ mutant, which makes unresected DSBs with Spo11 bound to the 5’ ends, thereby preventing repair by homologous recombination (Keeney *et al*. 1997; Neale *et al*. 2005). The Tel1 checkpoint kinase is recruited to the ends of the break, resulting in a low level of Mek1 kinase activation that is sufficient to delay exit from Prophase I but not prevent it completely (Carballo *et al*. 2008; Ho and Burgess 2011). When the MRC is abrogated, cells progress into Meiosis I with broken chromosomes, resulting in fragmented nuclei after Meiosis II (Figure 5C) (Mckee and Kleckner 1997). Similar to *NDT80-mid*, the *mek1-D78A E79A* exhibited fragmented nuclei and premature entry into Meiosis I compared to the *sae2*Δ diploid. The *mek1-D78A D79A* mutant also rescued the prophase I delay in the *hed*Δ*R* background.

### The FHA acidic loop is required for promoting proper Meiosis I chromosome segregation and spore viability

Two different proteins, Ndt80 and Rrm3, both of which are involved in either the regulation or execution of meiotic recombination, interact with Mek1 through a conserved RPxKR motif. The question then arises whether there might be additional Mek1 substrates that contain a similar sequence. Phenotypic analysis of the *mek1-AAA* and *mek1-D78A E79A* mutant suggest Ndt80 and Rrm3 are not the only proteins that interact with the acidic loop. The *mek1-D78A E79A* mutant is especially interesting in this regard. Two-hybrid experiments showed that the D78A E79A mutations weaken the interactions between *GAD-NDT80*/*GAD-RRM3* and *lexA-mek1-D78A E79A*, but do not eliminate them. However, in meiotic cells, *mek1-D78A E79A* exhibits as severe an MRC defect as *mek1*Δ in the *sae2*Δ background. Furthermore, when *mek1-D78A E79A* is combined with *hed*Δ*R*, spore viability is decreased to a far greater extent than *NDT80-mid* due to Meiosis I non-disjunction. We speculate that the Rad51-mediated recombination that occurs during meiosis in the *hed*Δ*R* background requires a higher level of Mek1 function to enable strand invasion into homologs, rather than sister chromatids. This idea is consistent with data showing the Dmc1 is better at making stable interhomolog connections than Rad51 (Callender *et al*. 2016). An exciting possibility is that the *mek1-D78A E79A* mutant encodes a kinase with a decreased affinity for an RPXKR protein that is phosphorylated by Mek1 to promote interhomolog bias.

## METHODS

### Yeast media and meiotic time courses

Yeast media and the sporulation protocol are described in (Lo and Hollingsworth 2011). For meiotic timecourses, sporulating cells were taken at different timepoints and assayed for a variety of parameters. Meiotic progression was measured by fixing cells with 3.7% formaldehyde, staining the DNA with 4’,6-diamidino-2-phenylindole (DAPI) and using fluorescence microscopy to count the number of nuclei. Red1 protein was detected using whole cell immunofluorescence with α-Red1 antibodies (Wan *et al*. 2004). Detailed protocols for each of these assays are described in (Ziesel *et al*. 2022). For immunoblot analyses, 5 mL sporulating cultures at each time point were pelleted in 15 mL conical tubes, the supernatants discarded and the pellets stored at −80°C.

### Yeast Strain Construction

Strains and their genotypes are shown in Table 1. All strains are derived from the SK1 background, except for L40 and DY1. For deletion of a gene, a DNA fragment containing the drug resistance marker *kanMX6*, *natMX4*, or *hphMX4* and 50 base pairs (bp) of homology flanking the open reading frame (ORF) of the gene to be deleted was amplified from the pFA6a-kanMX6, p4339, or pAG32 plasmids, respectively, by polymerase chain reactions (PCR) and transformed into the relevant yeast strains (Longtine *et al*. 1998; Goldstein and Mccusker 1999). To replace an endogenous gene promoter with the *CLB2* promoter (*P_CLB2_*), a DNA fragment containing *kanMX6* and *P_CLB2_* with 50 bp upstream of the gene of interest and downstream from *kanMX6* was amplified from pMJ787 by PCR and transformed into the relevant yeast strain.

Gene deletions were confirmed by PCR looking for the absence of the WT sequence and presence of the drug marker by combining a primer located upstream of the deleted ORF with either a gene-specific or drug-specific primer, respectively. Promoter replacements were confirmed using a forward primer located upstream of the inserted promoter and a reverse primer within the promoter.

For stable integration of various alleles, the *RRM3 URA3* plasmid, pBG22, or its mutant derivatives were digested with *Sna*BI for integration downstream of the *rrm3*Δ. The vector pRS306 was digested with NsiI for integration downstream of *ura3*. The *URA3 MEK1* integrating plasmid, pLP37, or its mutant derivatives, was digested with BamHI to target integration 800 bp upstream of the *MEK1* start codon. The presence of the plasmid borne alleles was verified by PCR using an upstream forward primer and an internal gene-specific reverse primer.

RCEN refers to diploids containing the *RFP* and *CFP* genes under the control of a spore autonomous promoter integrated at allelic positions adjacent to the centromeres of chromosome VIII (Thacker *et al*. 2011; Ziesel *et al*. 2022). The NH2740 RCEN and NH2783 RCEN diploids were created by deleting *MEK1* from the haploid parents of the NH2598 RCEN and NH2616 RCEN, respectively (Ziesel *et al*. 2022).

The *ndt80-mid sae2*Δ diploid (NH2759::pNH317^2^) was constructed by first deleting *NDT80* using *natMX4* from the haploid parents of NH1054. The *ndt80*Δ *sae2*Δ haploids were then transformed with pNH317 digested with SnaBI to target integration of the plasmid upstream of *ndt80*Δ. The haploids were then mated to make the diploid. For the *mek1*Δ *sae2*Δ diploid, NH2758, *MEK1* was replaced with *natMX4* in the haploid parents of NH1054 and the haploids then mated to make the diploid. The *mek1*Δ *ndt80*Δ diploid, NH2750, was created by sequentially deleting *NDT80* and *MEK1* from S2683 and RKY1145 and then mating the haploids to make the diploid.

### Plasmid Construction

Plasmid genotypes and sources are listed in Table 2. All mutant alleles were sequenced in their entirety to ensure the absence of unwanted mutations either by the Stony Brook University Sequencing Facility or by Plasmidsaurus (https://www.plasmidsaurus.com/).

To generate pJW1, an *RRM3^51-723^* PCR fragment was amplified from SK1 genomic DNA and cloned into pACTII, pre-digested with NcoI and EcoRI, to be in frame with the pre-existing *GAD* gene using the NEBuilder® HiFi DNA Assembly Master Mix (hereafter referred to as Gibson assembly) (New England BiolabsCat. # E2621). The KR>DD and KR>AA mutations changed K188 R189 codons (AAGAGG) within the *RRM3* ORF to aspartate (GATGAT) or alanine (GCGGCG), respectively. These mutations were introduced into pJW1 by site-directed mutagenesis using the QuikChange II kit (Agilent Technologies, Cat #200523) to make pJW1-KR>DD and pJW1-KR>AA. pJW2 was constructed similarly to pJW1, but two PCR fragments, *RRM3^51-184^* and *RRM3^190-723^*, were inserted into pACTII to delete the RPAKR coding sequence.

All of the *RRM3* mutants were derived from the *RRM3 URA3* integrating plasmid, pBG22 (Ziesel *et al*. 2022). The *RRM3-*Δ*RPAKR* (*RRM3-mid*) allele contained in pBG22-ΔRPAKR was constructed by Gibson Assembly cloning of two inserts amplified from pBG22 using primers containing 1overlapping homology lacking the RPAKR codons into EcoRI/ClaI-digested pRS306. The pBG22-ΔRPAKR-9A plasmid was made similarly, except the template for the PCR reactions was pBG22-9A. To introduce alanine or aspartic acid mutations at codons T37, T73, S85, S116, S125, S155, S165, S227, and S272, 717 bp fragments containing these mutations were synthesized by Azenta Life Sciences (pUC57-RRM3-9A and pUC57-RRM3-9D). pBG22-9A and pBG22-9D were constructed by Gibson Assembly cloning of three inserts into EcoRI-ClaI-digested pRS306: (1) the first insert stretches from 901 bp upstream of the *RRM3* ORF to bp 49 within the ORF and was amplified from pBG22, (2) the second insert stretches from bp 49 to bp 866 in the *RRM3* ORF and was amplified from pUC57-RRM3-9A or pUC57-RRM3-9D, (3) the third insert stretches from bp 866 in the *RRM3* ORF to 378 bp downstream of the ORF and was amplified from pBG22. To make the pBG22-3A, 3D and 3E plasmids that contain *rrm3-3A*, *rrm3-3D* and *rrm3-3E* respectively, 1497 bp fragments containing the indicated mutations in codons T565 T692 T721 were synthesized by Azenta Life Sciences. These fragments begin 1334 bp upstream of the *RRM3* stop codon and end 163 downstream of the stop codon. There is an endogenous NheI site within the ORF and an endogenous SnaBI site within the 3’ untranslated region (UTR). The plasmids were digested with NheI and SnaBI and the resulting 1.5 kb fragments were isolated and ligated into NheI/SnaBI digested pBG22.

The *rrm3-12D* allele was similarly constructed by cloning the 1.5 kb NheI/SnaBI fragment from the Azenta 3D plasmid into Nhe1/SnaBI cut pBG22-9D. This approach could not be used to make the *rrm3-12A* allele, because one of the 9A mutations coincidentally created a second NheI site within the *RRM3* ORF. The pBG22-12A plasmid was therefore created by Gibson assembly of two PCR fragments into EcoRI/ClaI digested pRS306. One PCR fragment contained homology to the EcoRI site in pRS306, the 5’ UTR and the first half of the *rrm3-9A* ORF, while the other fragment contained homology to the ClaI site in pRS306, the second half of the *rrm3-3A* ORF and the 3’ UTR.

The plasmids, pGAD-SRS2^1035-1174^, pA8 and pA46 were isolated in a two-hybrid screen using *lexA-MEK1* as bait as described in (Chen *et al*. 2018). The codons encoding the RKSKR sequence in *GAD-SRS2^1035-1174^* were deleted using site directed mutagenesis to generate pGAD-SRS2^1035-1174^-ΔRKSKR. The *lexA-mek1* mutants shown in Figure 4 and Figure S1 were generated by site directed mutagenesis of pTS3. The plasmids used for the two-hybrid experiments shown in Figure S4 were constructed as follows. The polylinker in the B42 activation domain (B42AD) containing vector, pJG4-5, was extended by insertion of a sequence between the EcoRI and XhoI sites of the polylinker to create pJG4-5EP (for extended polylinker). The additional sequence is in frame with B42AD-HA (Figure S4A). The *B42AD-HA-NDT80^287-627^*gene fusion was made using PCR to amplify a fragment from DY-1 genomic DNA encoding amino acids 287-627 of *NDT80* flanked by ApaI and XhoI restriction enzyme sites. The ApaI site in the *NDT80* fragment was engineered to be in frame with the ApaI site located just downstream of *HA* in pJG4-5EP. The ApaI-XhoI digested *NDT80* fragment was ligated into ApaI/XhoI cut pJG4-5EP to make pJG4-5EP NDT80ΔDBD. The *MEK1* gene was similarly cloned into pJG4-5EP, except using EcoRI and BglII as the flanking restriction sites. Site directed mutagenesis was performed on the pJG4-5EP MEK1 plasmid to introduce the *R51A*, *M84A Y86A* and *W75A* mutations into *MEK1*. Fragments containing *MEK1* or *mek1* mutant alleles were isolated from these by plasmids after digestion with EcoRI and XhoI. These fragments were then subcloned into EcoR-XhoI digested pEG202, to fuse the *mek1* coding sequences in frame with *lexA*.

Complementation tests were performed using derivatives of the *URA3 MEK1* integrating plasmid, pLP37. With one exception, all of the mutations were introduced by site directed mutagenesis. The exception was pLW208, which contains the *mek1-AAA* allele. To make this plasmid, pLP37 was digested with BfuAI and AflII and two vector backbone BfuAI/AflII fragments (2.5 and 2.9 kb) were ligated to a 1.3 kb BfuAI/AflII fragment isolated from pTS3-AAA.

### Yeast two-hybrid assays

For the two-hybrid experiments shown in Figures 1, 4 and Figure S1, plasmids encoding *TRP1 lexA* or *LEU2 GAD* fusions were co-transformed into L40 by selection on SD-trp-leu medium. L40 contains two reporter genes, *HIS3* and *lacZ*, downstream of four *lexA* operators (Hollenberg *et al*. 1995). Protein-protein interactions were detected by (1) the ability to grow on SD-trp-leu-his medium and (2) the production of β-galactosidase. For spotting assays, transformants were grown in SD-leu-trp liquid medium at 30°C overnight. Culture samples were diluted 1:10 and the optical density at 660 nm (OD_660_) was spectrophotometrically determined. To ensure that the same number of cells were assayed for each transformant, culture volumes equal to two OD_660_ units were pelleted, and the cells resuspended in 100 *µ*l sterilized water. Cells were serial diluted ten-fold and 10 *µ*l undiluted and diluted cells were spotted onto SD-Leu-Trp or SD-Leu-Trp-His plates. The plates were incubated at 30°C for three days. β-galactosidase activity was detected using a colorimetric assay using 5-bromo-4-chloro-3-indolyl β-D-galactopyranoside (X-gal). Four *µ*l undiluted cells were spotted onto filter paper placed on the agar surface of an SD-leu-trp plate. After one day at 30°C, the filter paper was dipped into liquid nitrogen and laid on top of a second filter paper soaked with 0.03% X-gal in dimethyl formamide, Reactions were left at 30°C until completion for several hours (Chen *et al*. 2018).

For the two-hybrid experiments shown in Figure S4, the DY-1 yeast strain was transformed with pSH18-34 which carries the *lacZ* gene with *lexA* operators in the promoter, as well as derivatives of the *lexA* fusion plasmid, pEG202 and the B42 activation domain fusion (*B42AD*) plasmid, pJG4-5EP. Liquid β-galactosidase assays were performed as described in (Ausubel *et al*. 1994).

### BLAST Analysis

The full-length sequence of Rrm3 (YHR031C; SGD ID S000001073) from the *Saccharomyces cerevisiae* SK1 background was pasted into NCBI’s protein BLAST (Blastp) tool (https://blast.ncbi.nlm.nih.gov/Blast.cgi) and searched against the “non-redundant protein sequences with a word size of 6 and using BLOSUM62 substitution matrix. The first 100 hits were filtered for non-redundant alignments (by species) and ones containing gaps in the subject sequence that were aligned to the query’s RPAKR (amino acid 185-189) were filtered out. The remaining alignments were uploaded to http://weblogo.threeplusone.com/ to generate a logogram.

### AlphaFold Multimer analysis

Structural predictions were generated using AlphaFold Multimer (Evans *et al*. 2022). Predictions were run using an interface available at Google Colab (https://colab.research.google.com/github/deepmind/alphafold/blob/main/notebooks/AlphaFold.ipynb).

### *In vitro* Mek1 kinase assays

GST-Mek1 was partially purified from meiotic yeast cells as described in (Lo and Hollingsworth 2011). GST-Mek1-as, purified 3XFLAG-Rrm3, 6-Fu-ATP*γ*S (10 mM; BIOLOG Life Science Institute F008), and 10 mM ATP were thawed on ice. 25 *µ*l kinase reactions contained GST1-Mek1-as, 3XFLAG-Rrm3 or BSA, and 4% dimethylsulfoxide (DMSO) in kinase buffer (50 mM Tris-HCl (pH7.5), 200 mM NaCl, 10 mM MgCl_2_), with 4 *µ*M ATP and 400 *µ*M 6-Fu-ATP*γ*S added last to initiate the reaction. For analog-inhibited reactions, a 10 mM stock of 1-(1,1-dimethylethyl)-3-(1-naphthalenyl)-1*H*-pyrazolo[3,4-*d*]pyrimidin-4-amine (1-NA-PP1; Cayman Chemical 10954) in DMSO was thawed on ice and added to a final concentration of 15 *µ*M. Reactions were incubated at 30°C for 30 minutes and then thiophosphates were alkylated by the addition of 2.5 mM *p*-nitrobenzyl mesylate (PNBM; Abcam ab138910) followed by incubating at room temperature for 2 hours (Lo and Hollingsworth 2011). 26 *µ*l of 2x Laemmli sodium dodecyl sulfate (SDS) sample buffer (100 mM Tris-HCl (pH 6.8), 4% SDS, 10% /J-mercaptoethanol, 6% glycerol, 0.2% bromophenol blue) were added and the samples were heated at 95°C for 5 minutes before storage at −20°C. 20 *µ*l of the kinase reaction were used for immunoblot analysis using the a-thiophosphate ester antibody (Abcam ab133473).

### Protein Preparation and Western Blotting

Yeast culture samples (15×10^7^ cells for meiotic cultures) were spun down and pellets were frozen at −80°C. The thawed pellets were resuspended in 5 mL 5% trichloroacetic acid (TCA) and inverted at 4°C for 10 minutes to allow protein precipitation. The TCA-treated pellets were vigorously washed in 1 mL acetone once by high-intensity vortexing before acetone was removed and the pellets fully air-dried. The dried pellets were resuspended in 200 *µ*l of lysis buffer (50 mM Tris-HCl pH 7.5, 1 mM EDTA, 2.75 mM DTT, 1.1 mM PMSF, Roche protease inhibitor at 2x working concentration (Roche 04693132001)), beaten with ∼200 *µ*l 0.5 mm glass beads (Biospec 11079105) using a FASTprep machine (MP Biochemicals) at 6 m/sec for 40 seconds. 150 *µ*l of 2x Laemmli SDS sample buffer was added to the lysate and boiled at 95°C for 5 minutes before pelleting and collecting the supernatant. Prepared protein samples were stored at −20°C until use.

Protein samples were loaded onto a 7.5% or 12% polyacrylamide gels (Bio-Rad Cat #3459902) and run at 150 to 200 Volts in running buffer (25 mM Tris, 0.1% SDS, 190 mM glycine) until appropriate fractionation was achieved. Proteins were transferred onto a polyvinylidene difluoride (PVDF) membrane that was first treated by successive soaking in ethanol, water, and transfer buffer (250 mM Tris, 200 mM glycine). The membrane was then incubated in blocking buffer (5% (w/v) dry milk powder dissolved in TBST (250 mM NaCl, 20 mM Tris-HCl (pH 7.5), 0.1% Tween-20)) for at least 30 minutes. Primary antibodies were added to the blocking buffer at the dilutions indicated in Table S3 and membranes rocked for 4 hr at room temperature or overnight at 4°C. Blots were washed three times with TBST for 5 minutes each with rocking, then incubated in blocking buffer containing horse radish peroxidase (HRP) bound secondary antibody and washed again as before. Proteins were detected by chemiluminescence using a luminol-H_2_O_2_ solution (WesternBright ECL Kit, K-12045-D20). Membranes were were in contact with the luminol-H_2_O_2_ solution for 2 minutes and then exposed for imaging using the Chemidoc imaging system (Bio-Rad).

### Southern Blot Analysis

At the indicated timepoints, 10 ml sporulating culture was added to 10 ml 95% ethanol and frozen at −20°C. Genomic DNA was isolated from the thawed cell pellets using the MasterPure Yeast DNA Purification kit (Lucigen MPY80200). The DNA was digested with XhoI and NgoMIV and fractionated on a 0.6% agarose gel. After the DNA was transferred by capillary action to Hybond-N+ membrane (Amersham #RPN303B), the membrane was probed and processed as described in (Owens *et al*. 2018). The blots were imaged using a Fuji Phosphoimager FLA7000 with the Multi-Gauge Analysis Software. TIFF files were processed using ImageJ with the maximum contrast limit of contrast set at 1748.

### Antibodies

The sources and dilutions used for various antibodies are listed in Table S3. Rrm3 polyclonal antibodies were generated by Covance Research Products (now Labcorp Drug Development) in a rabbit (NY2524) using the peptide, NERVKDFYKRLETLK (aa 709-723), located at the very C-terminus of the protein as antigen. The antibodies were affinity purified using the same peptide by the company. The specificity of the antibody was verified by the detection of a protein with approximately the predicted molecular weight (79.5 kD) in protein extracts from vegetatively grown *RRM3*, but not *rrm3*Δ, strains (Figure 2B).

## Data Availability Statement

All of the raw data used for the figures in this paper can be found in the Supplemental Material (Data S1). Statistical analyses were performed using GraphPad Prism 9.0 for Mac.

## ACKNOWLEDGEMENTS

Many thanks to Teresa de los Santos who did the two-hybrid screen many years ago that identified various *lexA-MEK1* interacting *GAD* fusion genes. We are grateful to members of the Hollingsworth lab for helpful discussion and Andrew Ziesel and Bob Gaglione for help with plasmid/strain constructions. Bob Gaglione and Raunak Dutta provided helpful comments on the manuscript.

## Supplemental Figure legends

**Figure S1. Two-hybrid assays using various *lexA-MEK1* mutants.** (A) *lexA-MEK1* plasmids encoding single amino acid substitutions in the acidic loop (pTS3-D78A, pTS3-E79A and pTS3-D80A) were tested for two-hybrid interactions with various *GAD* fusions as described in Figure 4. The *lexA-MEK1* control panel is the same as in Figure 4. (B) Immunoblot of protein extracts from the strains used in (A) probed with either α-Mek1 or α-Arp7 antibodies. The asterisk indicates a non-specific band. The “V” and “WT” lanes are the same as in Figure 4. (C) *lexA-mek1* plasmids encoding either lysine (pTS3-E333K) or alanine (pTS3-E333A) substituted for E333 in the Mek1 activation loop were tested for two-hybrid interactions with various *GAD* fusions as described in Figure 4. (D) Immunoblot of protein extracts from the strains used in (C) probed with either Mek1 or Arp7 antibodies.

**Figure S2. The Mek1 interaction motif in Rrm3 is predicted by AlphaFold Multimer to interact with an acidic patch in the Mek1 FHA domain, as well as a glutamate in the Mek1 activation loop.** (A) Structural model generated by AlphaFold Multimer showing the interaction between a 44 amino acid sequence from Rrm3 (aa 161-205) containing the RPAKR Mek1 interaction motif with full-length Mek1. The color code is the same as in Figure 3. Green and yellow indicate the Mek1 FHA domain with yellow showing the b-sheets and intervening loop containing amino acids that were mutated for phenotypic characterization. Cyan indicates the Rrm3 peptide. Orange indicates E333 in the Mek1 activation loop. (B) Close-up of the region indicated by the box in Panel A showing the side chains of D78 E79 D80 and E333 in Mek1 relative the R185, P186, A187, K188 and R189 side chains of Rrm3.

**Figure S3. Confidence levels of the AlphaFold-predicted structures of Mek1-Ndt80 and Mek1-Rrm3.** Confidence levels are represented using the predicted local-distance difference test (pLDDT) metric (Jumper *et al*. 2021). The pLDDT values are per-residue estimated confidence levels scaled from 0-100. The values were converted using a coloring scheme and plotted onto the modeled structures using ChimeraX. (A) Mek1-Ndt80. Box 1 and Box 2 indicate the regions shown in Figure 3. (B) Mek1-Rrm3. The box indicates the region shown in Figure S2.

**Figure S4. Amino acids in the lateral surface of a β-sheet in the Mek1 FHA domain are required for interaction with Ndt80 in the two-hybrid system.** (A) The sequence added to the polylinker of pJG4-5 to make pJG4-5EP. (B) The yeast strain, DY-1, carrying a *lacZ* reporter gene on the 2μ *URA3* plasmid, pSH18-34, was transformed with *B42AD-NDT80^287-627^* (pJG4-5EP-Ndt80ΔDBD) and one of the following plasmids: *lexA* (pEG202-indicated by “-“), *lexA-MEK1* (pEG202 MEK1), *lexA-mek1-R51A* (pEG202 mek1-R51A), *lexA-mek1 M84A Y86A* (pEG202 MY), or *lexA-mek1 W73A V75A* (pEG202 WV). Transformants were assayed for two-hybrid interactions by measuring b-galactosidase activity in liquid cultures as described in (Gyuris *et al*. 1993; Duncker *et al*. 2002). Each dot represents a biological replicate. Error bars indicate the means and standard deviations. (C) Immunoblot of protein extracts from the strains shown in A. Ponceau S staining was used as a protein loading control. A single blot was simultaneously probed with lexA and HA antibodies to detect lexA-Mek1 and B42AD-HA-NDT80*^287-627^*, respectively. The blot was then incubated with an α-mouse secondary antibody conjugated to AlexaFluor 488 to detect HA and an α-rabbit secondary antibody conjugated to AlexaFluor 647 to detect lexA.

---

**Table S1.**
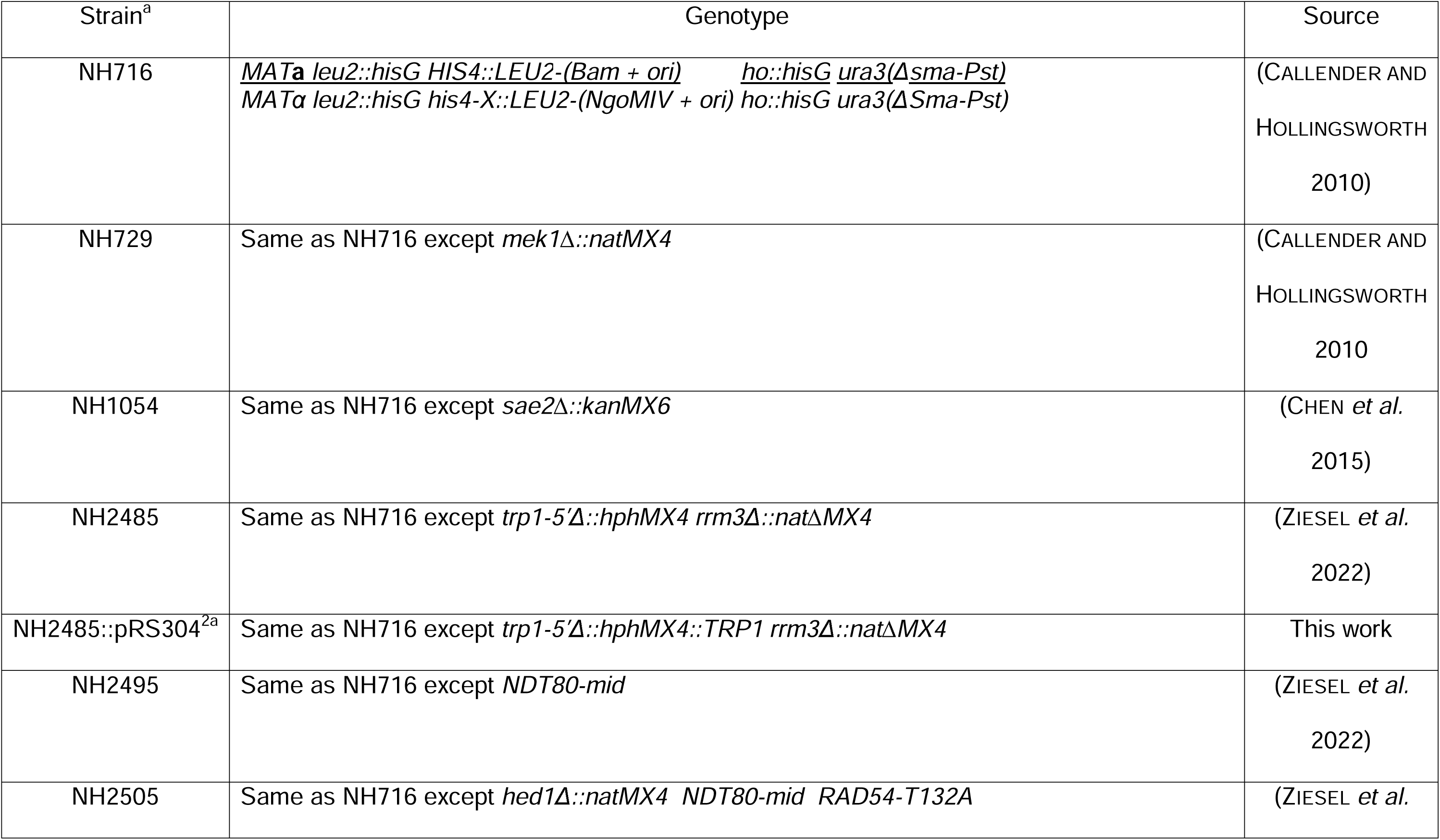
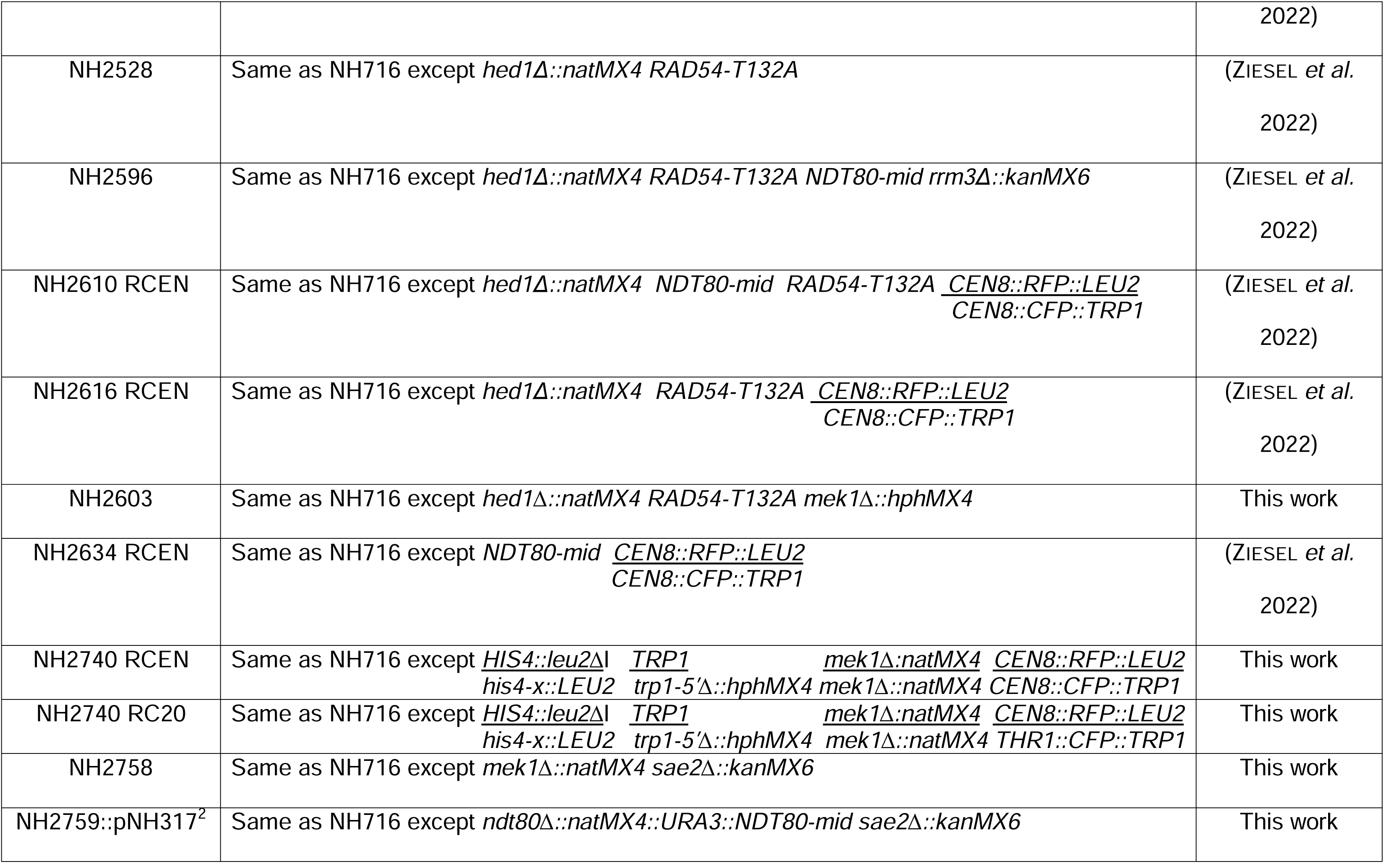
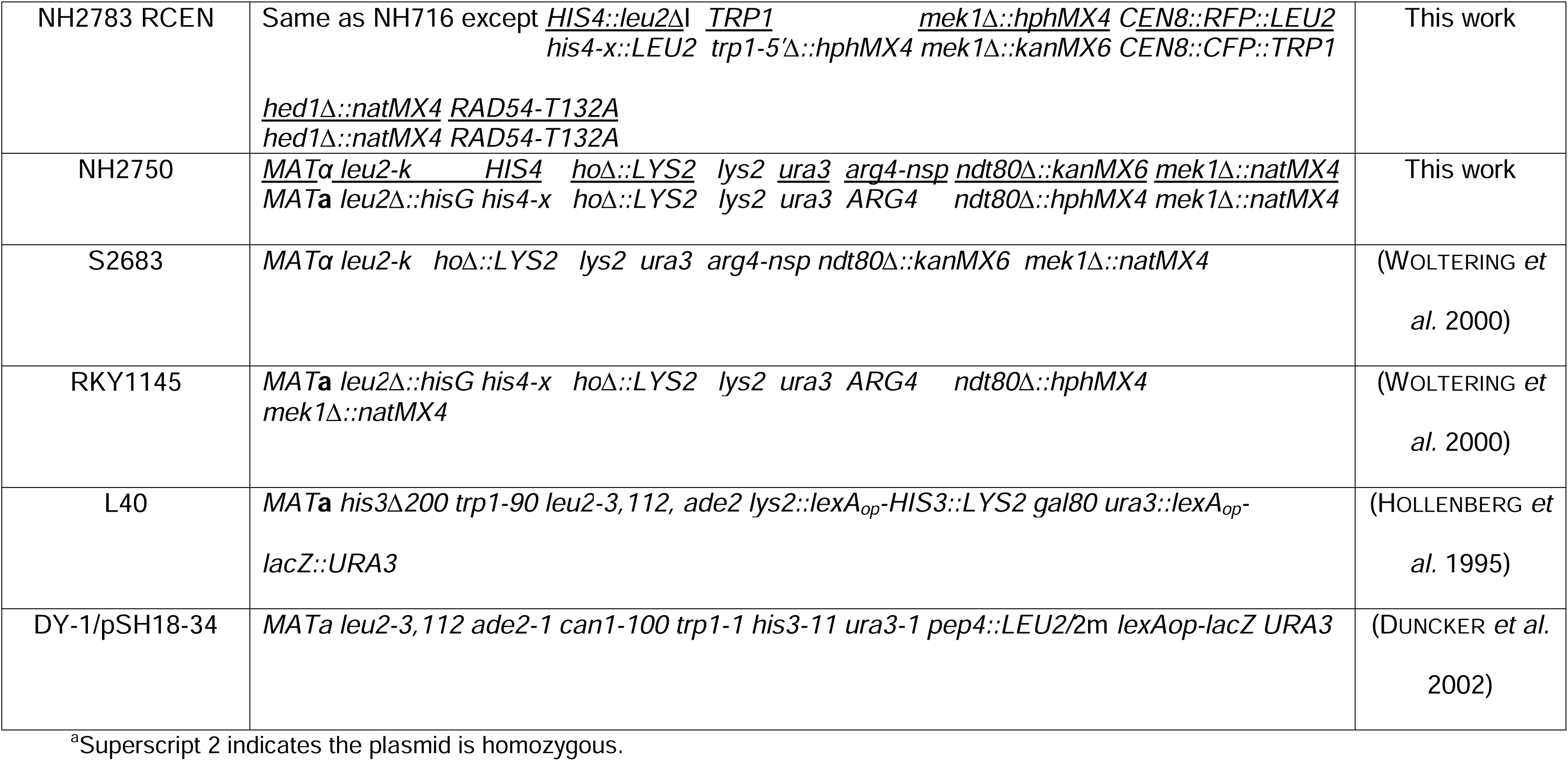
*Saccharomyces cerevisiae* strains.

**Table S2.**
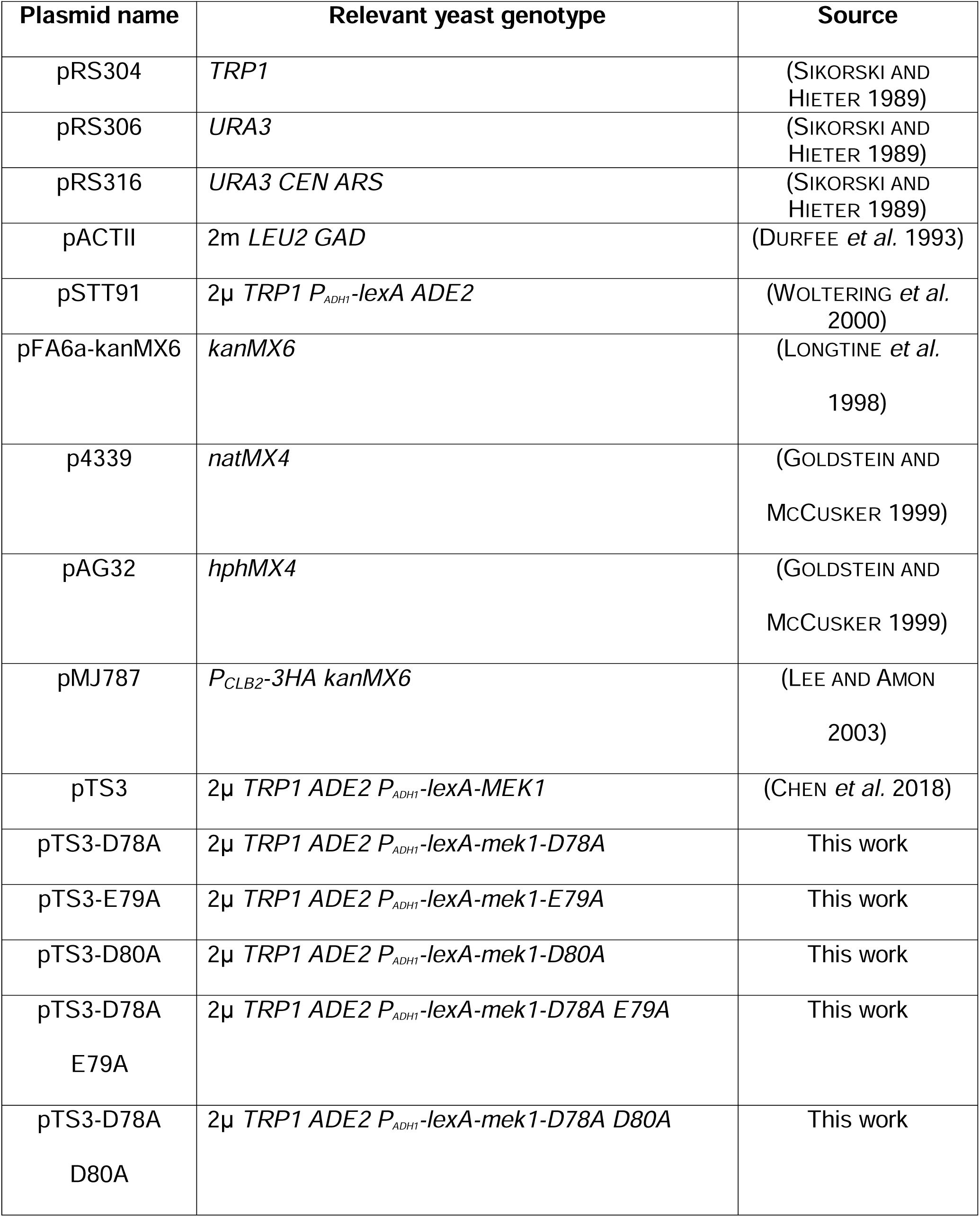
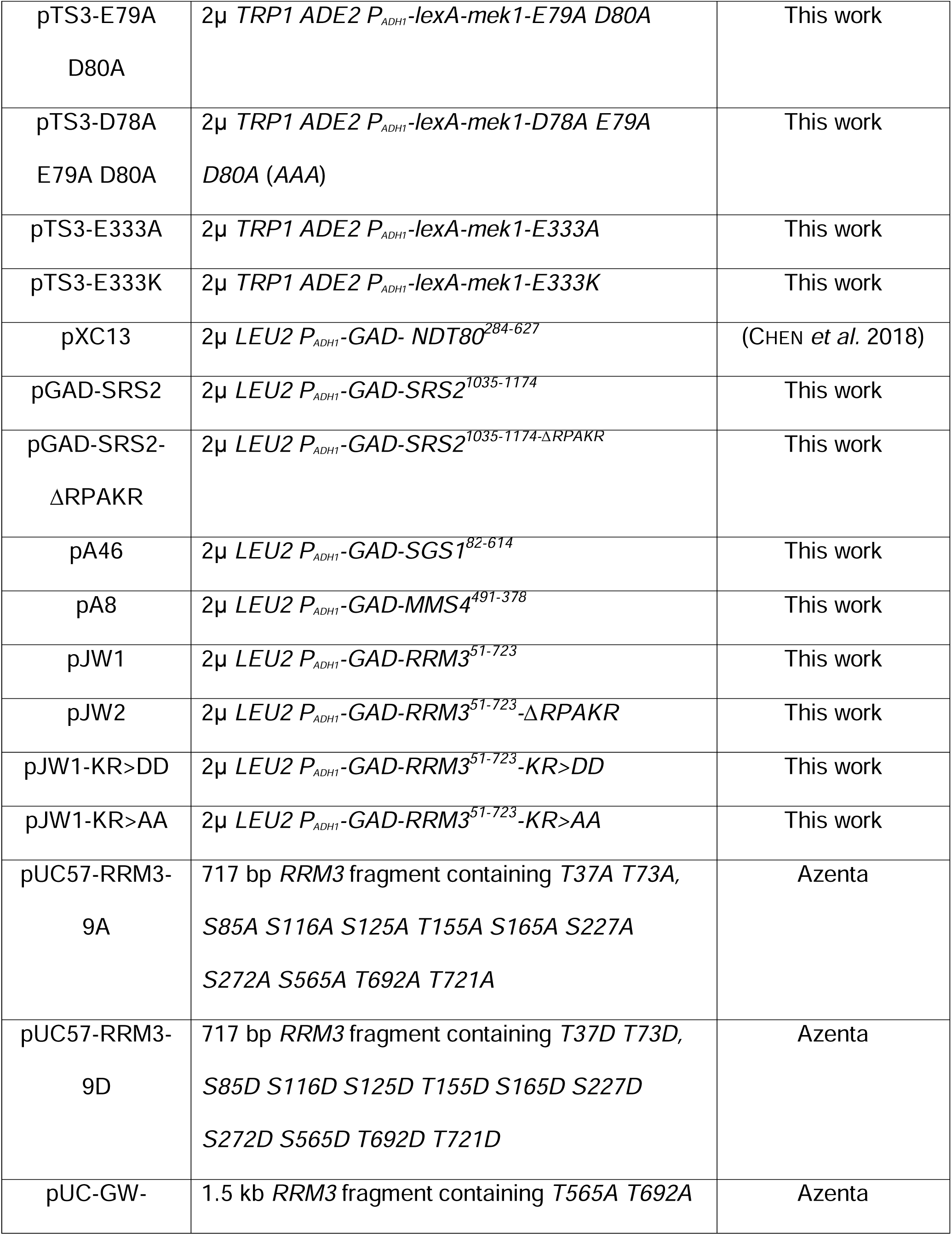
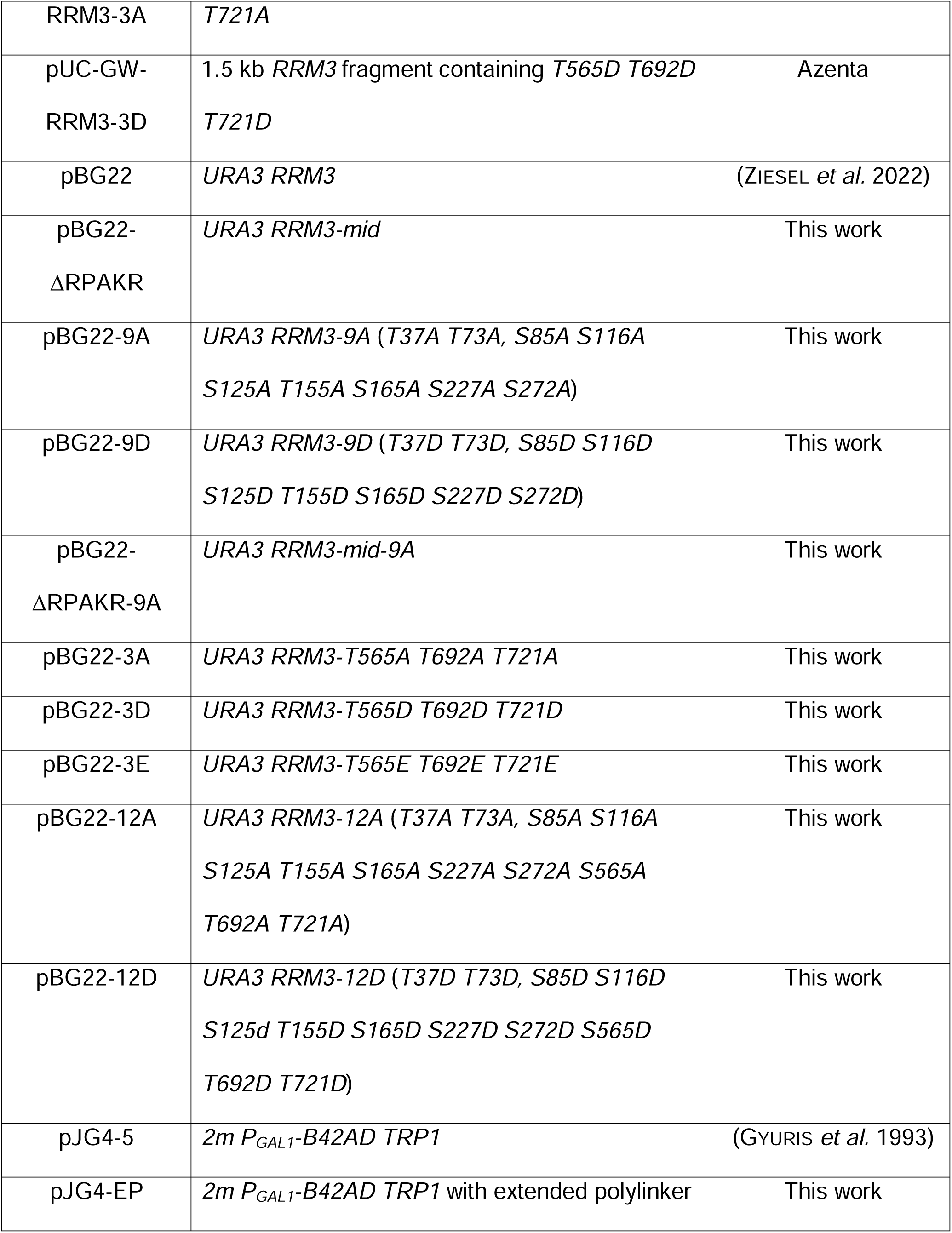
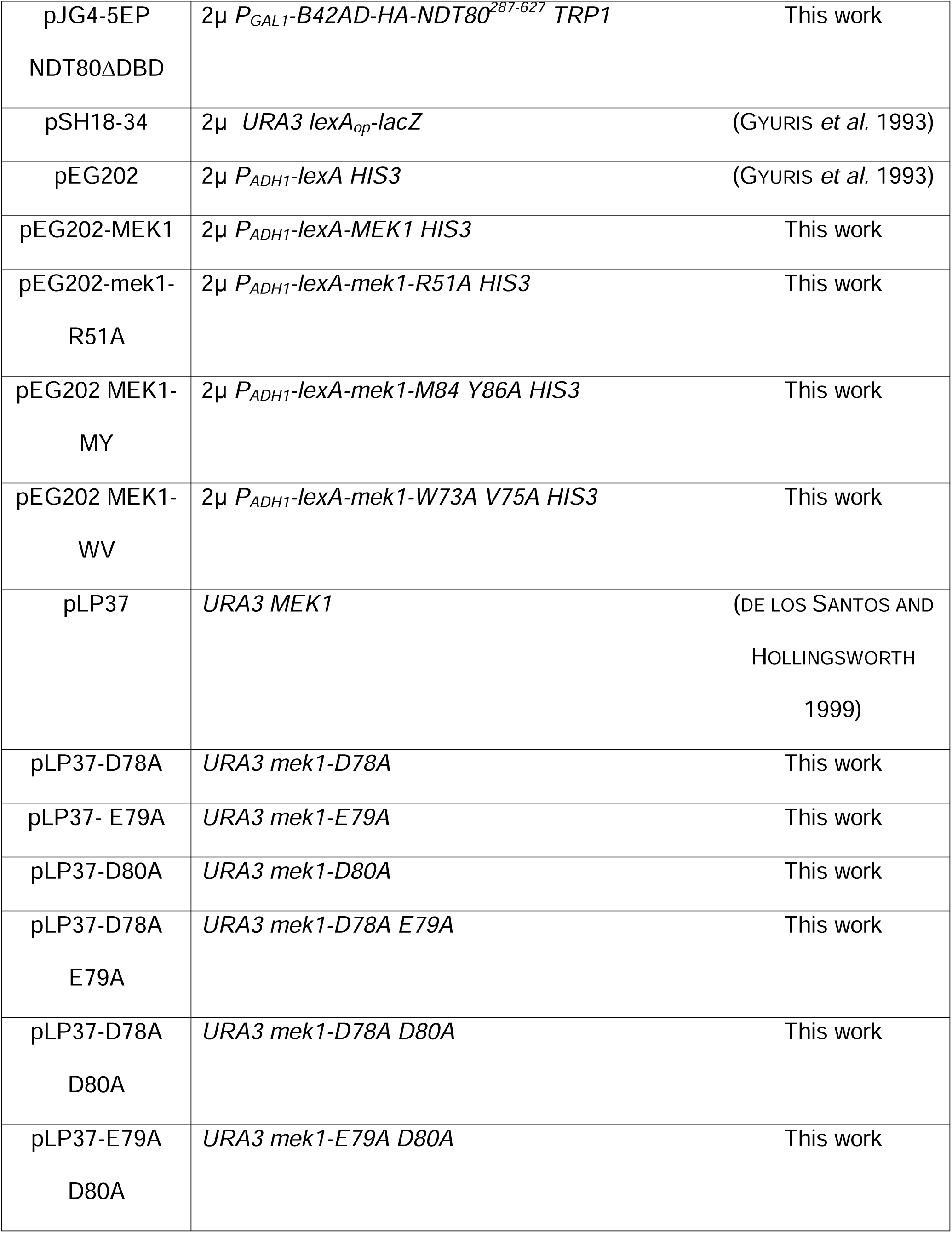
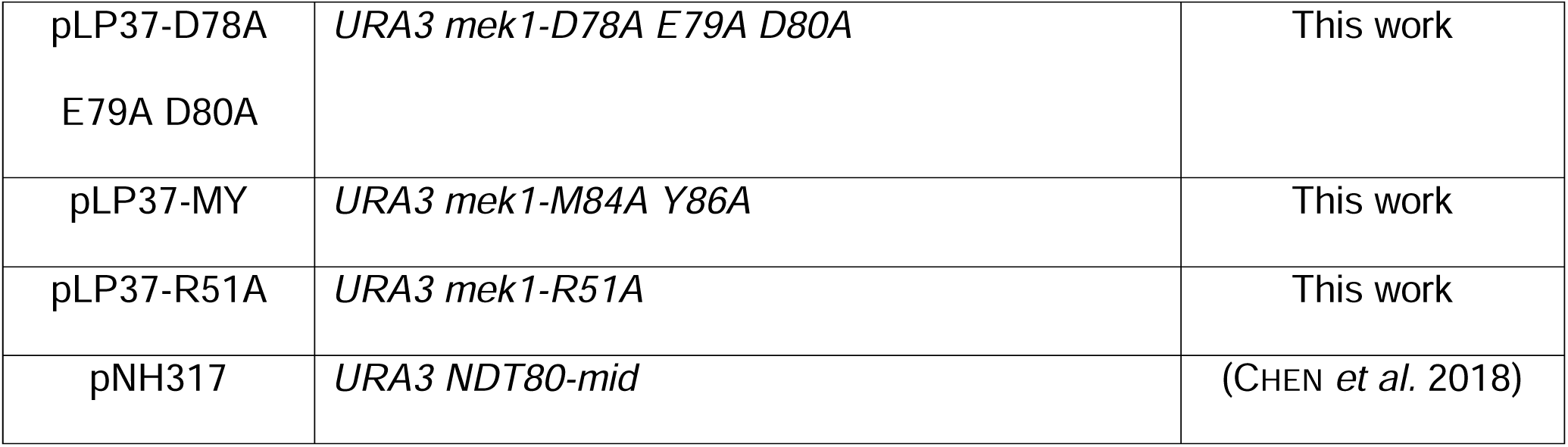
Plasmids.

**Table S3.** Antibodies.

| Protein | Primary (1<sup>o</sup>) | Dilution | Source (cat #) | Secondary (2<sup>o</sup>) | Dilution | Source (catalog #) |
| --- | --- | --- | --- | --- | --- | --- |
| Arp7 | Goat α-Arp7 | 1:5,000 | Santa Cruz (sc-8961) | Mouse α-goat | 1:10,000 | Santa Cruz (sc-2354) |
| GAD | Mouse α-GAD | 1:10,000 | Takara (630402) | Goat α-mouse | 1:10,000 | Invitrogen (PI32430) |
| Mek1 | Guinea pig α-Mek1 (NY1970) | 1:5,000-1:20,000 | (PRUGAR *et al.* 2017) | Goat α-Guinea pig | 1:5,000-1:10,000 | Jackson Immuno research (106-035-003) |
| Mek1-pT327 | Phospho-S/T Akt substrate | 1:10,000 | Cell Signalling (9611S) | Goat α-rabbit | 1:10,000 | Invitrogen (31460) |
| Rrm3 | Rabbit α-Rrm3 | 1:10,000 | This work | Goat α-rabbit | 1:10,000 | Invitrogen (31460) |
| phosphosites | Rabbit Thio-phosphate ester | 1:5000 | Abcam ab133473 | Goat α-rabbit | 1:10,000 | Invitrogen (31460) |
| Red1 | Red1 | 1:300 | (WAN *et al.* 2004) | Goat α-rabbit Alexa 488 | 1:1000 | Invitrogen (A11008) |

