## Supplementary figures and images for "An acidic loop in the FHA domain of the yeast meiosis-specific kinase Mek1 interacts with a specific motif in a subset of Mek1 substrates"

### Figure S1

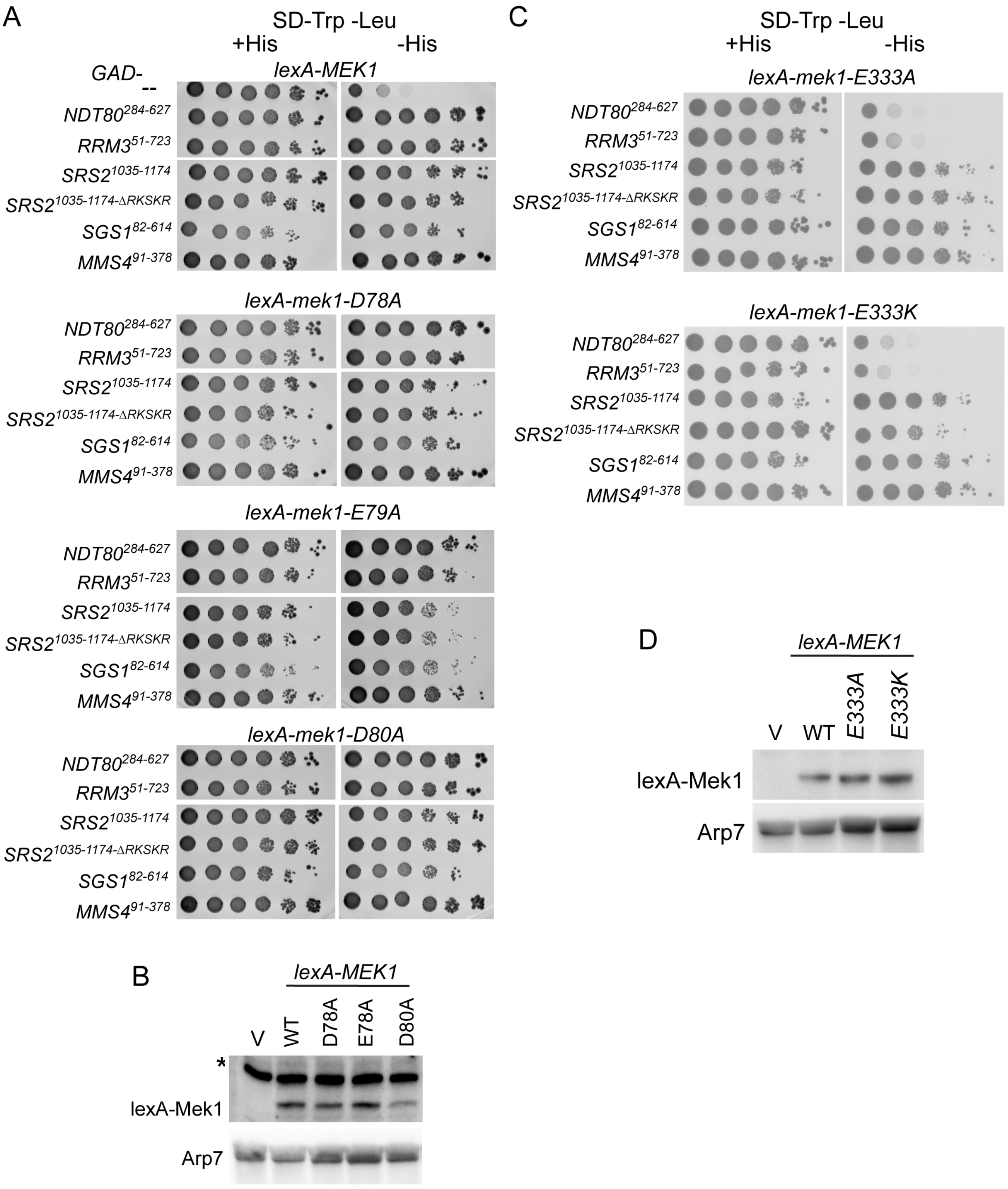

### Figure S2

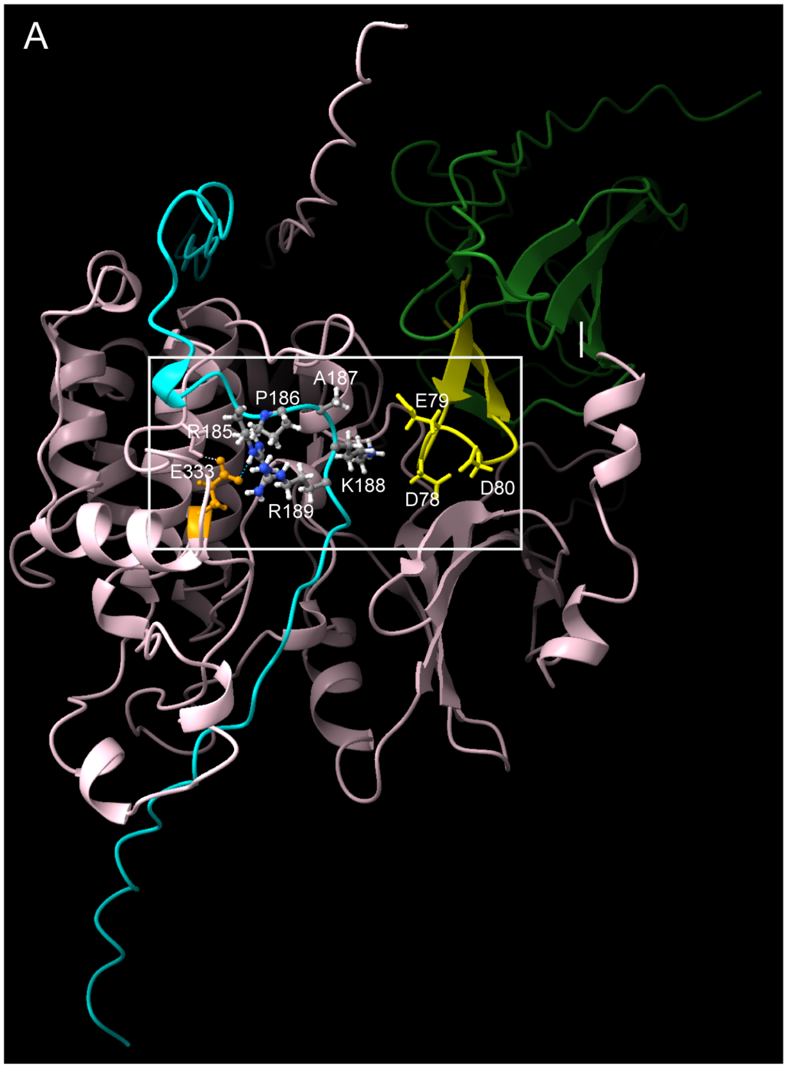

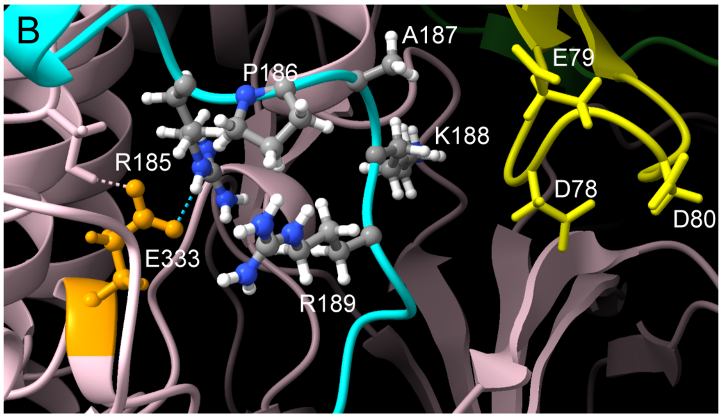

### Figure S3

FigureS3

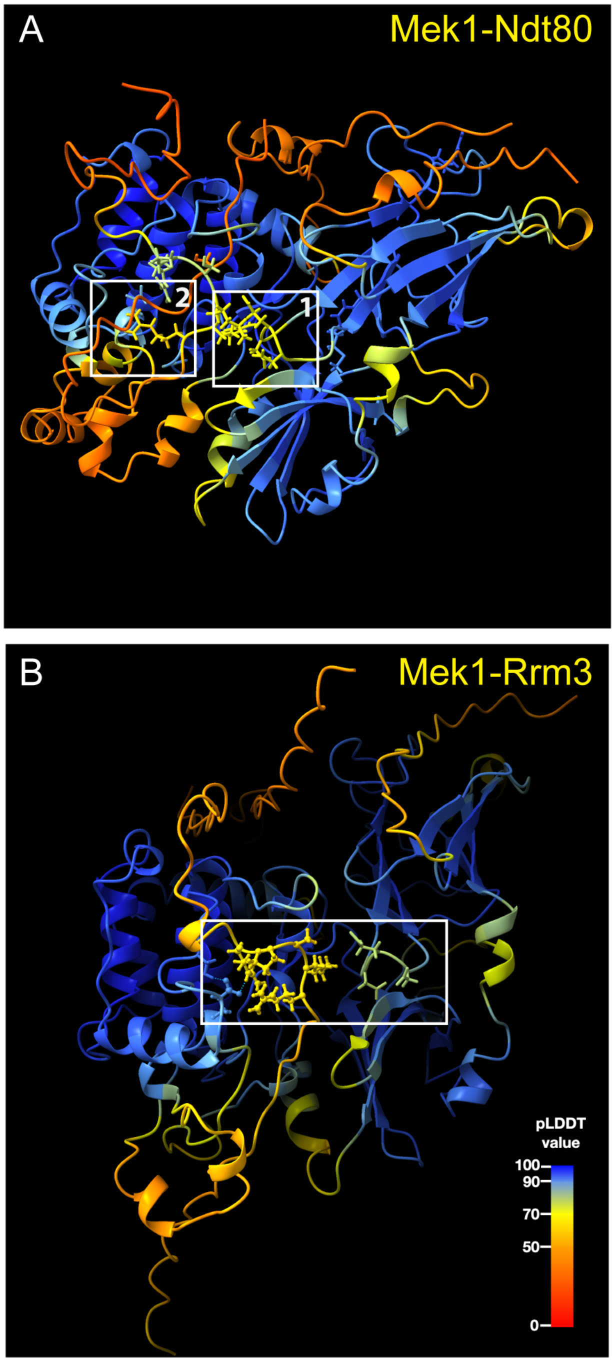
